# The N-glycosylation defect in Lec5 and Lec9 CHO cells is caused by absence of the DHRSX gene

**DOI:** 10.1101/2024.06.18.599300

**Authors:** Takfarinas Kentache, Charlotte R. Althoff, Francesco Caligiore, Erika Souche, Céline Schulz, Julie Graff, Eline Pieters, Pamela Stanley, Joseph N. Contessa, Emile Van Schaftingen, Gert Matthijs, François Foulquier, Guido T. Bommer, Matthew P. Wilson

## Abstract

Glycosylation-deficient Chinese hamster ovary (CHO) cell lines have been instrumental in the discovery of N-glycosylation machinery. Yet, the molecular causes of the glycosylation defects in the Lec5 and Lec9 mutants have been elusive, even though for both cell lines a defect in dolichol formation from polyprenol was previously established. We recently found that dolichol synthesis from polyprenol occurs in three steps consisting of the conversion of polyprenol to polyprenal by DHRSX, the reduction of polyprenal to dolichal by SRD5A3 and the reduction of dolichal to dolichol, again by DHRSX.

This led us to investigate defective dolichol synthesis in Lec5 and Lec9 cells. Both cell lines showed increased levels of polyprenol and its derivatives, concomitant with decreased levels of dolichol and derivatives, but no change in polyprenal levels, suggesting DHRSX deficiency. Accordingly, N-glycan synthesis and changes in polyisoprenoid levels were corrected by complementation with human DHRSX but not with SRD5A3. Furthermore, the typical polyprenol dehydrogenase and dolichal reductase activities of DHRSX were absent in membrane preparations derived from Lec5 and Lec9 cells, while the reduction of polyprenal to dolichal, catalyzed by SRD5A3, was unaffected. Long-read whole genome sequencing of Lec5 and Lec9 cells did not reveal mutations in the ORF of *SRD5A3*, but the genomic region containing *DHRSX* was absent. Lastly, we established the sequence of Chinese hamster DHRSX and validated that this protein has similar kinetic properties to the human enzyme. Our work therefore identifies the basis of the dolichol synthesis defect in CHO Lec5 and Lec9 cells.

## Introduction

Glycosylation mutants affecting numerous proteins involved in the glycosylation process have been isolated in CHO cells mainly by selection for resistance to toxic lectins. More than 40 different complementation groups have been characterized in this way, and in many cases, mutations in glycosylation pathway genes have been identified (1, 2) However, the genetic defects in Lec5 (or B211) and Lec9 mutants have not been defined. These cell lines are characterized by reduced N-glycan synthesis and an alteration in the abundance of specific N-glycan subtypes, due to a defect in the conversion of polyprenol to dolichol (3–8).

Dolichol plays an important role in various types of glycosylation as a carrier of monosaccharides (Dol-P-glucose and Dol-P-mannose) and of the lipid-linked oligosaccharide (LLO) that is transferred *en bloc* onto the N-X-S/T/C motif of proteins (where N is asparagine, X is any amino acid except proline and S/T/C is serine, threonine or, rarely, cysteine) during N-glycosylation. Polyprenol, the precursor of dolichol, is formed via the mevalonate pathway and differs from dolichol only by the presence of a double bond between carbons 2 and 3 of the terminal isoprene unit (9). When polyprenol is used as a mono- or oligosaccharide carrier, this difference considerably alters the activity of several enzymes that usually act on different dolichol derivatives in the context of glycosylation (10–14).

Though the lack of conversion of polyprenol to dolichol in Lec5 and Lec9 cells is well established, the mechanism of this defect is still unknown (6, 8, 15, 16). Polyprenol conversion to dolichol was, in 2010, proposed to involve an NADPH-dependent reductase (17). The demonstration of the role of the SRD5A3 gene product in the conversion of polyprenol to dolichol led to the assumption that SRD5A3 was the long sought polyprenol reductase. Surprisingly, this finding did not lead to the identification of the defect in Lec5 and Lec9 cells, including in our hands (unpublished Sanger sequencing data).

The widely held view that SRD5A3 is a polyprenol reductase has been challenged recently by our group (18). We demonstrated that the conversion of polyprenol to dolichol is a three-step process, involving polyprenal and dolichal as intermediates, and in which SRD5A3 is a polypren*<u>a</u>*l (not a polypren*<u>o</u>*l) reductase. Another enzyme, encoded by the *DHRSX* gene, catalyzes the two other steps, *i.e.* the oxidation of polyprenol to polyprenal and the final reduction of dolichal to dolichol (**Fig. 1A**). In this work we show that the *DHRSX* gene is deleted in Lec5 and Lec9 cells, and that both their dolichol synthesis and N-glycan defects are rescued by human DHSRX.

**Figure 1.**
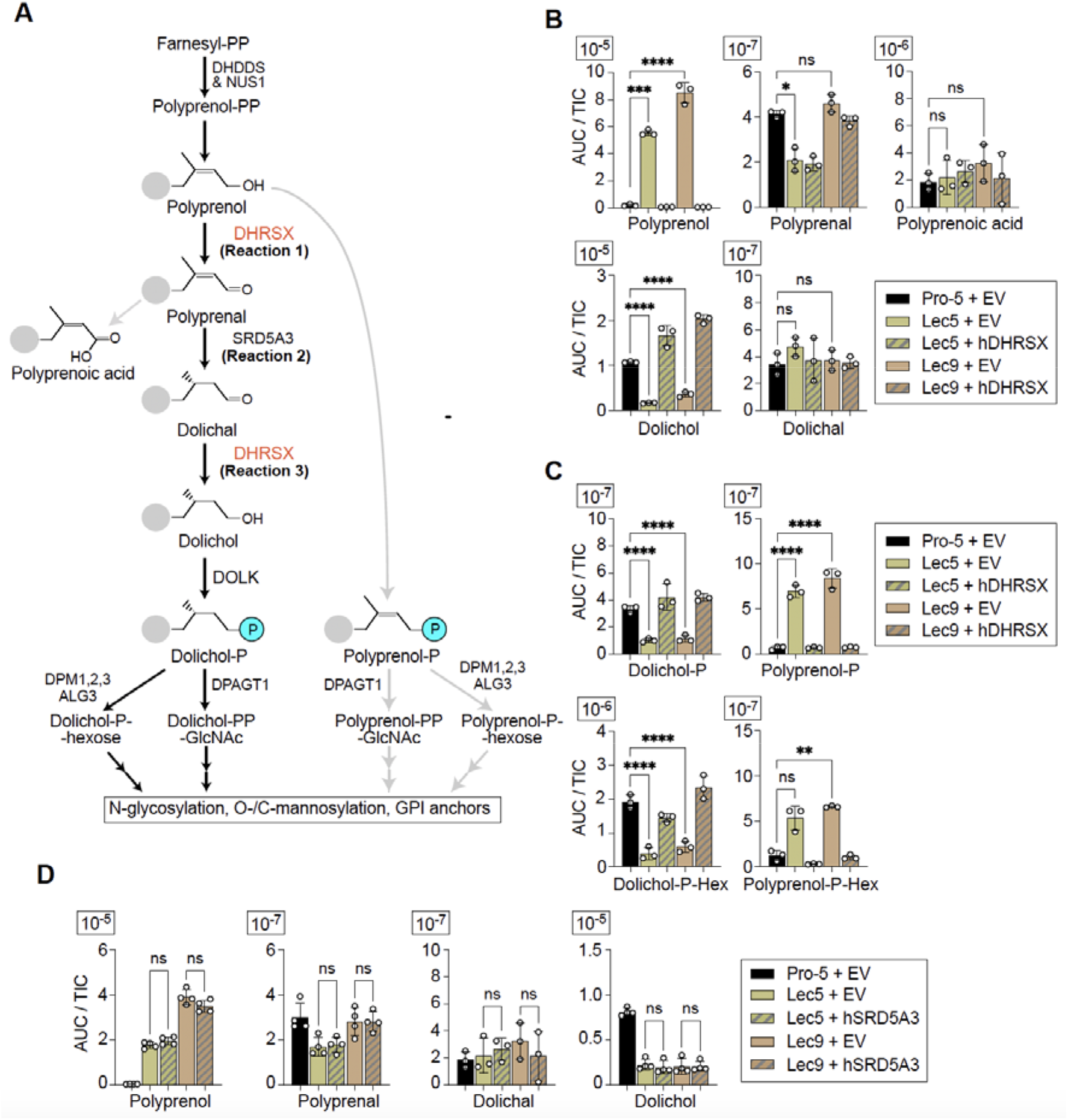
**Accumulation of polyisoprenoid species in Lec5/9 cells resembles that in *H. sapiens* DHRSX deficiency.** (A) Mammalian dolichol biosynthesis pathway. (B) Polyisoprenoid species in Pro-5 (control), Lec5 and Lec9 CHO cells complemented or not with human DHRSX. See **Fig. S1A** for all isoprenyl chain lengths. (C) Phosphoisoprenoid and hexose phosphoisoprenoid species in Pro-5 (control), Lec5 and Lec9 CHO cells complemented or not with human DHRSX. See **Fig. S1B** for all isoprenyl chain lengths. (D) Polyisoprenoid species in Pro-5 (control), Lec5 and Lec9 CHO cells complemented or not with human SRD5A3. See **Fig. S1C** for all isoprenyl chain lengths. B-D: Data represent area under the curve (AUC) normalized to total ion count (TIC) of species with 19 isoprenyl units (means ± SEM; n=3). For statistical analysis an ordinary one-way ANOVA followed by a Tukey’s multiple comparisons test was used; ns, **,*** and **** represent not significant, p < 0.01, p < 0.001, and p < 0.0001, respectively.

## Results

Accumulation of polyisoprenoid species in Lec5 and Lec9 cells resembles that in *H. sapiens* DHRSX deficiency.

Studies on human cell lines show that DHRSX deficiency leads to a decrease in dolichol and an increase in polyprenol, and similar changes in glycosylation to those previously observed with Lec5 and Lec9 cells (18). In particular, previous work on the glycosylation defect in CHO Lec5 and Lec9 cells showed that the conversion of polyprenol to dolichol is deficient (16, 19). As a result of this, N-glycan synthesis in Lec5 and Lec9 cells utilizes polyprenol-phosphate rather than dolichol-phosphate species, leading to perturbation in the synthesis and transfer of the LLO (**Fig. 1A**) (18).

These changes resemble those we found when inactivating SRD5A3 or DHRSX in human cell lines. In both cases we observed a strong accumulation of polyprenol-derived lipids alongside a reduced level in dolichol-derived lipids. Biochemically, the main feature that distinguishes DHRSX from SRD5A3 deficient cells is that only the latter accumulate the intermediate polyprenal and its carboxyl derivative, polyprenoic acid (**Fig. 1A**) (18).

To identify the defect causing Lec5 and Lec9 phenotypes, we analyzed isoprenoids in these cells compared to their parental cell line, Pro-5. As previously observed for SRD5A3- and DHRSX-deficient cells, we observed accumulation of polyprenol (28- and 43-fold for Lec5 and Lec9, respectively), polyprenol-phosphate (11- and 13-fold), and polyprenol-phosphohexose (4- and 5-fold) and a decrease in dolichol (6- and 3-fold), dolichol-phosphate (both by 3-fold), and dolichol-phosphohexose (5- and 3-fold) (**Fig. 1B-C**, see **Fig. S1A** for all measured lengths; 17 to 20).

However, Lec5 and Lec9 cells did not show a significant accumulation of polyprenal and polyprenoic acid, as is commonly observed in SRD5A3 deficiency. This suggested that the defective enzyme was DHRSX. Accordingly, complementation of Lec5 or Lec9 cells with *H. sapiens* (h) DHRSX restored polyisoprenoid species levels similar to those observed in control Pro-5 cells, whereas complementation with *H. sapiens* (h) SRD5A3 had no effect (**Fig. 1B-D**, see **Fig. S1A-B** for all measured lengths; 17 to 20).

### DHRSX activity is absent in Lec5 and Lec9 cells, but SRD5A3 activity is unaffected

DHRSX is a quite unique dehydrogenase/reductase in that it catalyzes two distinct reactions: i) the conversion of polyprenol to polyprenal (**Reaction 1**, **Fig. 1A**) and ii) that of dolichal to dolichol (**Reaction 3**, **Fig. 1A**). Accordingly, it also has similar affinities for NAD(H) and NADP(H). By contrast, SRD5A3 is a NADPH-specific reductase that converts polyprenal to dolichal (**Reaction 2**, **Fig. 1A**). These enzymatic activities can be readily measured in human cell extracts such as HAP1 and lymphoblastoid cells (18).

To confirm the identity of the enzymatic defect we assayed these activities in membrane preparations derived from Pro-5, Lec5 and Lec9 cells, including those expressing hDHRSX. Using polyprenol and NAD^+^ or NADP^+^ as substrates we clearly detected the formation of polyprenal in membranes derived from wild type cells, but not from Lec5 or Lec9 cells (**Fig. 2A**). As expected, expression of hDHRSX in Lec5 and Lec9 cells increased the formation of polyprenal even beyond that observed in Pro-5 cells.

**Figure 2.**
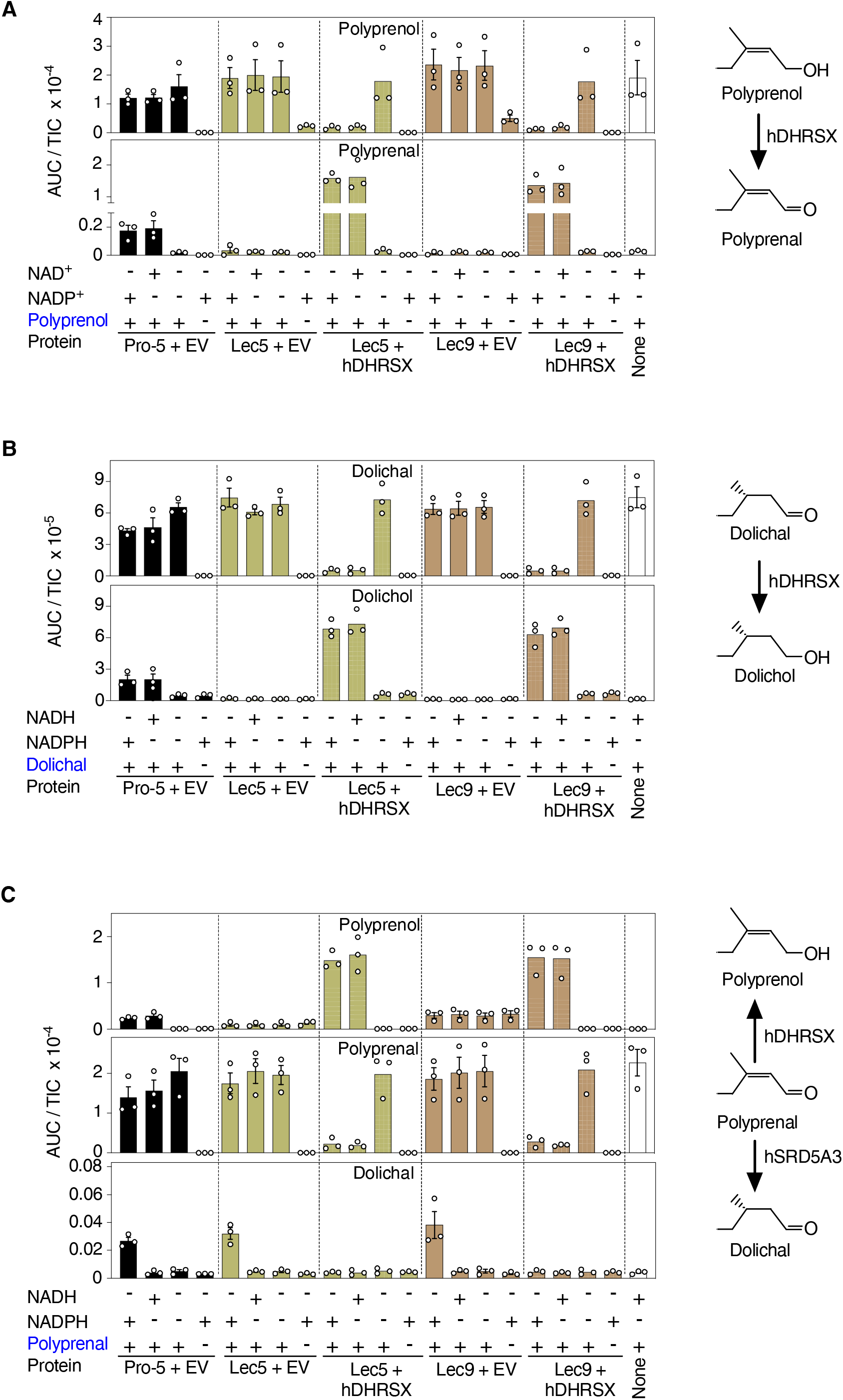
**DHRSX activites are absent in Lec5 and Lec9 cells, but SRD5A3 activity is unaffected** (A) Failure of Lec5 and Lec9 CHO cells to form polyprenal from polyprenol. Polyprenal formation from polyprenol was measured after incubation of 5 µg/mL of a 13 to 21 isoprene unit-containing polyprenol mixture with 5 mM NADP^+^ or NAD^+^ and 1 mg/mL membrane extracts for 2 h at 37 °C. Measurements are based on the formation of polyprenal with 18 isoprene units (means ± SEM, n= 3). Data is TIC-normalized AUC (means ± SEM, n=3). (B) Lec5 and Lec9 CHO cells lack dolichal reductase activity. Dolichal-18 and dolichol-18 were measured in reactions containing 1 mg/mL CHO membrane proteins, 5 μg/mL dolichal mixture, and 5 mM NAD(P)H for 2h at 37°C. Data is TIC-normalized AUC (means ± SEM, n=3). (C) Detection of NADPH-dependent polyprenal reductase activity mediated by SRD5A3 in Lec5 and Lec9 CHO cells. SRD5A3 activity was determined by measuring dolichal formation after incubation of 5 μg/mL of a polyprenal mixture with 5 mM NAD(P)H and 1 mg/mL membrane preparations for 2 hours at 37 °C. The reverse reaction of polyprenol dehydrogenase mediated by DHRSX (see Fig. 2A) was detected as well. The panel shows only species with 18 isoprene units (means +/- SEM; n=3). Data is TIC-normalized AUC (means ± SEM, n=3).

We also obtained similar results when we assessed the reduction of dolichal to dolichol, the second enzymatic function of DHRSX (**Fig. 2B**). Dolichol formation was clear in Pro-5 cells but absent in Lec5 and Lec9 cells. Again, expression of hDHRSX strongly increased dolichol formation. The absence of both activities characteristic for DHRSX indicates that the Chinese hamster ortholog of DHRSX is deficient in Lec5 and Lec9 cells, and that, similarly to the human enzyme, Chinese hamster (ch) DHRSX is able to catalyze both the conversion of polyprenol to polyprenal and the conversion of dolichal to dolichol, with no apparent specificity for NADP(H) or NAD(H).

To ensure that the SRD5A3 activity was not affected in Lec5 or Lec9 cells, we incubated the same cell membrane preparations with polyprenal and either NADH or NADPH. We noted a comparable formation of dolichal in Pro-5, Lec5 and Lec9 cells, in the presence of NADPH, but not with NADH. This indicated that SRD5A3 activity was normal in Lec5 and Lec9 cells (**Fig. 2C**).

Surprisingly, the formation of dolichal from polyprenal seemed to be abolished in cells overexpressing DHRSX. This is explained by the fact that DHRSX catalyzes, *in vitro,* the reduction of polyprenal to polyprenol in the presence of either NADH or NADPH, leading to the strong increase of polyprenol formation in membrane preparations from cell lines overexpressing DHRSX (**Fig. 2C**). Consequently, the substrate of SRD5A3, polyprenal, will be rapidly depleted leading to the formation of polyprenol at the expense of dolichal.

Taken together, these enzymatic assays indicated that DHRSX activity is absent in Lec5 and Lec9 cells, and that there is no deficiency of SRD5A3 in either cell line.

### The glycosylation defect in Lec5 and Lec9 cells is rescued by expression of human DHRSX

We used two complementary methods to assess glycosylation status in Lec5 and Lec9 cells: in depth characterization of an artificial glycosylation marker protein via western blotting, and an unbiased proteomic characterization of microsomal proteins.

Human DHRSX-deficient cell lines showed reduced glycosylation of LAMP2 (18). Unfortunately, antibodies directed at the human protein did not allow us to detect this protein in CHO cells (data not shown). We therefore used an artificial protein, Halo3N, containing three well-defined N-glycosylation sites. Halo3N is a derivative of the HaloTag® (20) engineered to contain three N-glycosylation sites, an EGFR signal sequence and a KDEL ER-retention signal, allowing the quantification of N-glycosylation site occupancy based on different migration profiles by SDS-PAGE analysis (**Fig. 3A**) (21).

**Figure 3.**
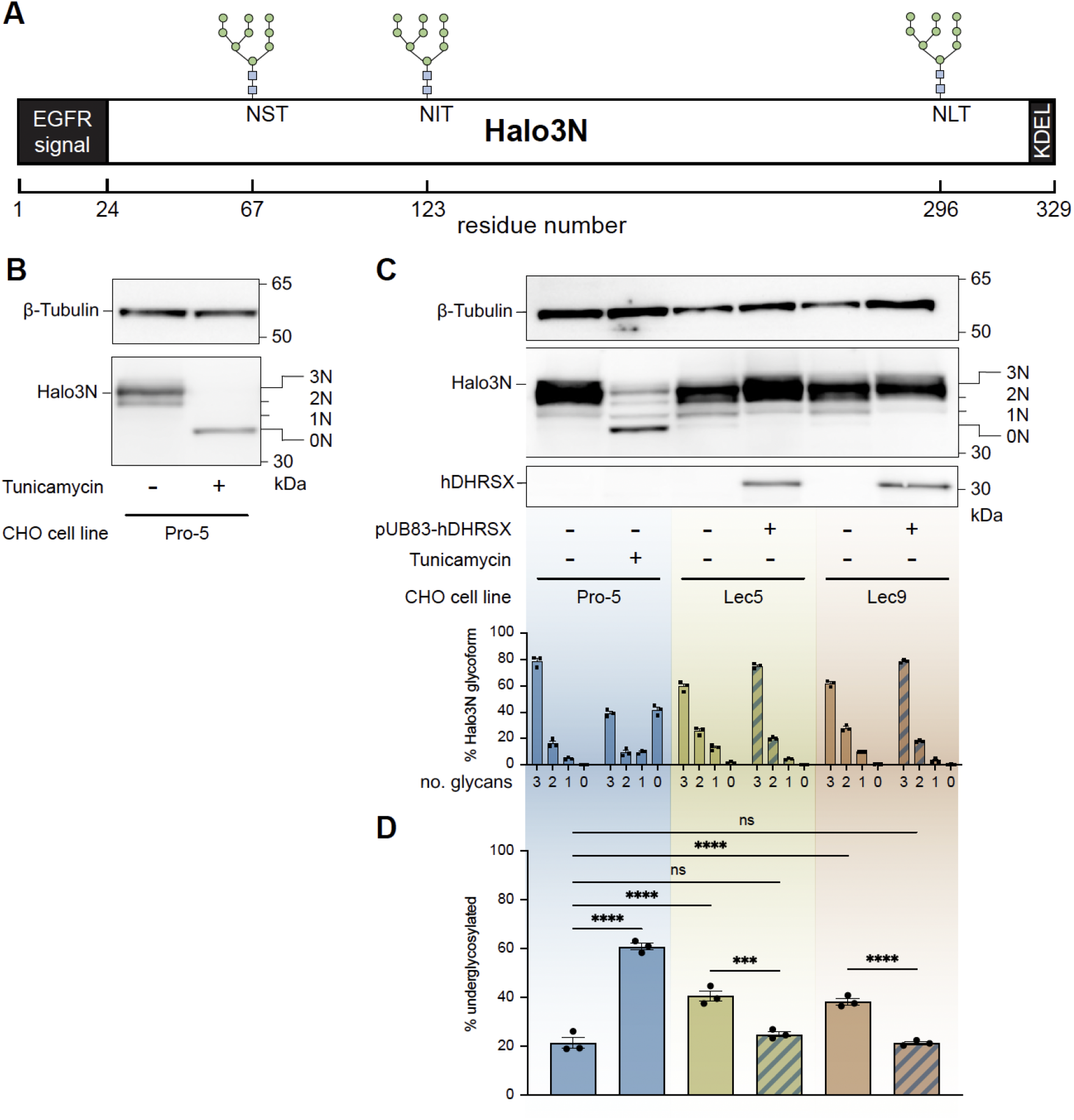
**The glycosylation defect in Lec5 and Lec9 cells is rescued by expression of human DHRSX** (A) The Halo3N glycomarker protein with N-glycosylation sites indicated. (B) Halo3N is a sensitive marker of N-glycosylation status in Pro-5 CHO cells. SDS-PAGE analysis shows that tunicamycin treatment (1μg/mL) for 24h abolishes all N-glycan attachment to Halo3N and signal derived from probing with a HaloTag® antibody is condensed to a single band (0N). Blot was also probed with a β-tubulin antibody as a protein loading control. (C) Lec5 and Lec9 CHO cells have deficient N-glycan attachment, rescued by expression of pUB83-hDHRSX. Probing with a HaloTag® antibody revealed that treatment with a proteasomal inhibitor (MG-132, 10 μmol/L, 24h) allows improved resolution of intermediate (2N, 1N and 0N) Halo3N glycosylation states in tunicamycin (1 μg/mL for 24h) treated Pro-5 cells. This facilitated their quantification in untreated Pro-5, Lec5 and Lec9 cells. hDHRSX expression was confirmed by reprobing with a DHRSX antibody and the blot was also probed with a β-tubulin antibody as a protein loading control (means +/- SEM; n=3). (D) Underglycosylation of the Halo3N reporter protein in Lec5 and Lec9 cells is restored to levels in Pro-5 control cells by expression of hDHRSX. % underglycosylated represents the signal intensity of the 0N, 1N and 2N glycosylation states combined, after probing with a HaloTag® antibody, compared to overall signal including the 3N glycosylation state. ns, *** and **** represent not significant, p < 0.001, and p < 0.0001, respectively. For statistical analysis an ordinary one-way ANOVA followed by a Tukey’s multiple comparisons test was used.

After stable expression of Halo3N in Pro-5, Lec5 and Lec9 cells, we first validated that the migration profile of this protein was indeed dependent on the glycosylation status in CHO cells. To this end, we analyzed lysates from Pro-5 cells cultured in the presence and absence of tunicamycin, an inhibitor of DPAGT1, the enzyme that transfers N-acetylglucosamine-1-phosphate onto dolichol phosphate in the early stages of lipid-linked oligosaccharide biosynthesis. Halo3N has four possible glycosylation states, carrying 3, 2, 1, or no N-glycans (hereafter 3N, 2N, 1N and 0N). Most of the signal in samples from untreated Pro-5 cells was concentrated in a single band, presumably corresponding to the fully glycosylated form (3N), whereas samples from Tunicamycin-treated Pro-5 cells showed a single band with a faster migration in SDS-PAGE, presumably corresponding to the unglycosylated form (0N; **Fig. 3B**).

Next, we compared Pro-5 with Lec5 and Lec9 cells and cell lines in which human DHRSX was overexpressed. These experiments were performed in the presence of the proteasome inhibitor MG-132 to limit potential degradation of hypoglycosylated proteins. This allowed us to detect four bands for Halo3N, presumably corresponding to the glycoforms 0N, 1N, 2N and 3N. Halo3N in control Pro-5 CHO cells was observed to be 79% fully glycosylated (3N), with 16% and 5% of the 2N and 1N forms, respectively (**Fig. 3C & 3D**). Both Lec5 and Lec9 cells showed a reduction in the fully glycosylated form and a concomitant significant increase in the hypoglycosylated forms. Both changes were rescued by expression of hDHRSX, leading to Halo3N glycosylation states undistinguishable from that of Pro-5 cells.

We also explored the glycosylation state of Pro-5, Lec5 and Lec9 cells via a proteomic approach. Inactivation of DHRSX in human HAP1 cells leads to a shift from dolichol to polyprenol as the LLO/monosaccharide carrier and to an increase in the transfer of immature glycans onto proteins (**Fig. 4A**) (18). We detected several Hexose(Hex)-3, Hex-4, Hex-5, and even Hex-6 glycans in Lec5 and Lec9 cells, which were absent in Pro-5 cells or when we re-expressed hDHRSX (**Fig. 4B, Table S1**). Extension of the linear Man-5 glycan is much less efficient when the LLO is assembled on polyprenol pyrophosphate as opposed to dolichol pyrophosphate (**Fig. 4A**) (12). Thus, the Hex-5 peptides increased in Lec5 and Lec9 cells are likely the result of the transfer of linear Man-5 onto proteins.

**Figure 4.**
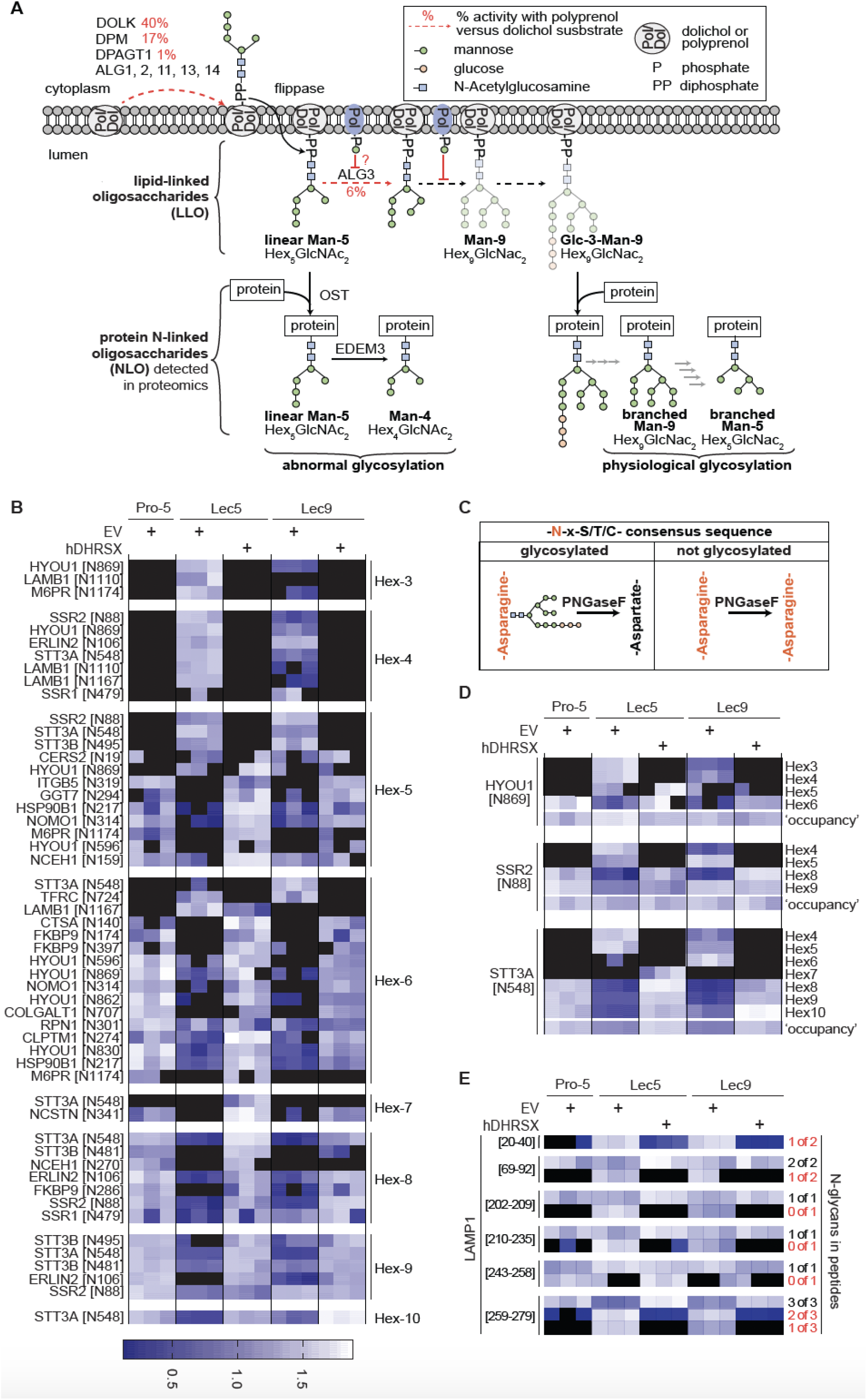
**Lec5 and Lec9 cells show glycan structures that disappear upon expression of human DHRSX.** (A) Schematic representation of the formation of N-linked glycans under physiological conditions and when dolichol is largely replaced by polyprenol (18). Note that the maturation beyond linear Man-5 LLO is inefficient in the latter situation. (B) Quantification of the indicated glycopeptides in membrane fractions of Pro-5, Lec5, and Lec9 cell lines. Each cell line was transduced either with a lentiviral vector containing an empty vector (EV), or, in case of Lec5 and Lec9, a vector inducing the expression of human DHRSX (hDHRSX). The heatmap represent the abundance different N-glycosylation sites, regrouped by glycosylation types from Hex-3 to Hex-10. Signals obtained in 3 biological replicates were log2-transformed and are shown relative to the 70^th^ percentile within each row to facilitate visualization. Black cells indicate missing values (see Supplementary Table S1 for non-normalized data). (C) Schematic of the PNGase F reaction on N-glycosylation sites. Hydrolytic removal of the attached glycans forms an aspartate residue, which can be distinguished from the asparagine residue present in the non-glycosylated peptide. (D) Detailed comparison of the different glycosylation states at selected glycosites, presented as in panel (B). This panel also includes data on the abundance of PNGase F-treated samples (‘occupancy’). (E) Abundance of peptides derived from the protein LAMP1 carrying 0, 1, 2 or 3 N-glycans. Peptides were quantified by nano-LC-MS/MS after treatment with PNGaseF, which converts glycan-carrying asparagine residues to aspartates. The numbers on the left indicate the location of the peptides within LAMP1. The numbers on the right indicate how many of the possible glycosylation sites were occupied. Hypoglycosylated forms are highlighted in red.

Linear Man-5 can be subjected to two enzymatic processes: On the one hand, a glucosylation/deglucosylation cycle plays a role in permitting proper protein folding. Accordingly, some of the Hex-6 glycans can be formed by transfer of glucose residues to Man-5 using ER glucosyltransferases, leading to Man-5/Glc-1 (22). On the other hand, demannosylation via the enzyme EDEM3 can target unfolded and hypoglycosylated proteins to ER-associated protein degradation (ERAD) (23). Accordingly, the Hex-4 and Hex-3 glycans appearing in Lec5 and Lec9 cells likely correspond to Man-4 and Man-3 glycans (21).

Conversely, a distinct subset of Hex-5 glycans and several Hex-6, Hex-7 and Hex-8 glycans were lost in Lec5 and Lec9 cells but reappeared upon expression of human DHRSX, likely representing intermediates in the physiological N-glycosylation pathway, including branched (as opposed to linear) Man-5 glycans. In Lec5 and Lec9 cells, the shift towards immature N-glycans at the expense of the fully extended N-glycans was also apparent when we analyzed individual peptides for which several different glycosylation states were observed (**Fig. 4D**). Even though comprehensive data are available only for a limited number of proteins, our findings document a clear change in glycan structures in Lec5 and Lec9 cells.

To explore whether these changes affect overall N-glycosylation site occupancy, we treated peptides with PNGaseF before proteomic analysis. This removes N-glycans, resulting in an aspartate residue which can be distinguished from the asparagine to which the glycan was attached (**Fig. 4C**). Changes in the occupancy were rather subtle or absent for the peptides presented in panels B and D (**Fig. 4C-D; ‘occupancy’**). Yet, when focusing on several peptides of the highly glycosylated protein LAMP1, we observed considerable heterogeneoity between glycosylation sites (**Fig. 4E**). Hypoglycosylated or unglycosylated forms appeared for all peptides, but only in one instance this was paralleled by a reduction in the abundance of the fully glycosylated form (amino acids 259 – 279, Fig. 4E). This suggested that, rather than N-glycans being absent altogether, in many cases mature glycans might be replaced by immature glycans, agreeing with previous glycoproteomic studies on human LLO biosynthesis enzyme deficiencies (24, 25).

Overall, we document a quantitative change evidenced by reduced Halo3N glycosylation and a site-dependent reduction in glycosylation site occupancy, as well as a structural change evidenced by the attachment of immature N-glycans to glycoproteins in Lec5 and Lec9 cells. Both these changes were rescued by expression of hDHRSX.

### CHO cells have a functional DHRSX ortholog that is absent in Lec5 and Lec9 CHO cells

A number of lectin-resistant CHO glycosylation mutant cell lines have either deletions or point mutations inactivating genes important for N-glycosylation (1). We therefore investigated whether a genetic mechanism was causing the DHRSX deficiency in Lec5 and Lec9 cells. Initially, attempts to determine the genomic and RNA-derived coding sequences of *DHRSX* via Sanger sequencing in Lec5/9 cells were unsuccessful. A partial *DHRSX* sequence was acquired in Pro-5 cells, but efforts were hampered by sequence differences, as compared to those of available *C. griseus* reference sequences (e.g. CHO-K1 CriGri_1.0, GenBank: GCA_000223135.1; CriGri-PICRH-1.0, GenBank: GCA_003668045.2; CHOK1S_HZDv1, GenBank: GCA_900186095.1).

To overcome this problem, we performed Oxford Nanopore Technology (ONT) long-read whole genome sequencing (WGS) using genomic DNA extracted from Pro-5, Lec5 and Lec9 CHO cells and performed *de novo* genome assembly for each cell line. This led to full-length assemblies that were of similar quality to those previously published (26) (**Table 1, Table S2**). In Pro-5 cells, *DHRSX* was detected towards the end of a contig of 385 kbp (contig 8156; **Fig. S2 & Fig. 5D**). We were unable to identify reads pertaining to *DHRSX* in Lec5 or Lec9 cells.

**Figure 5.**
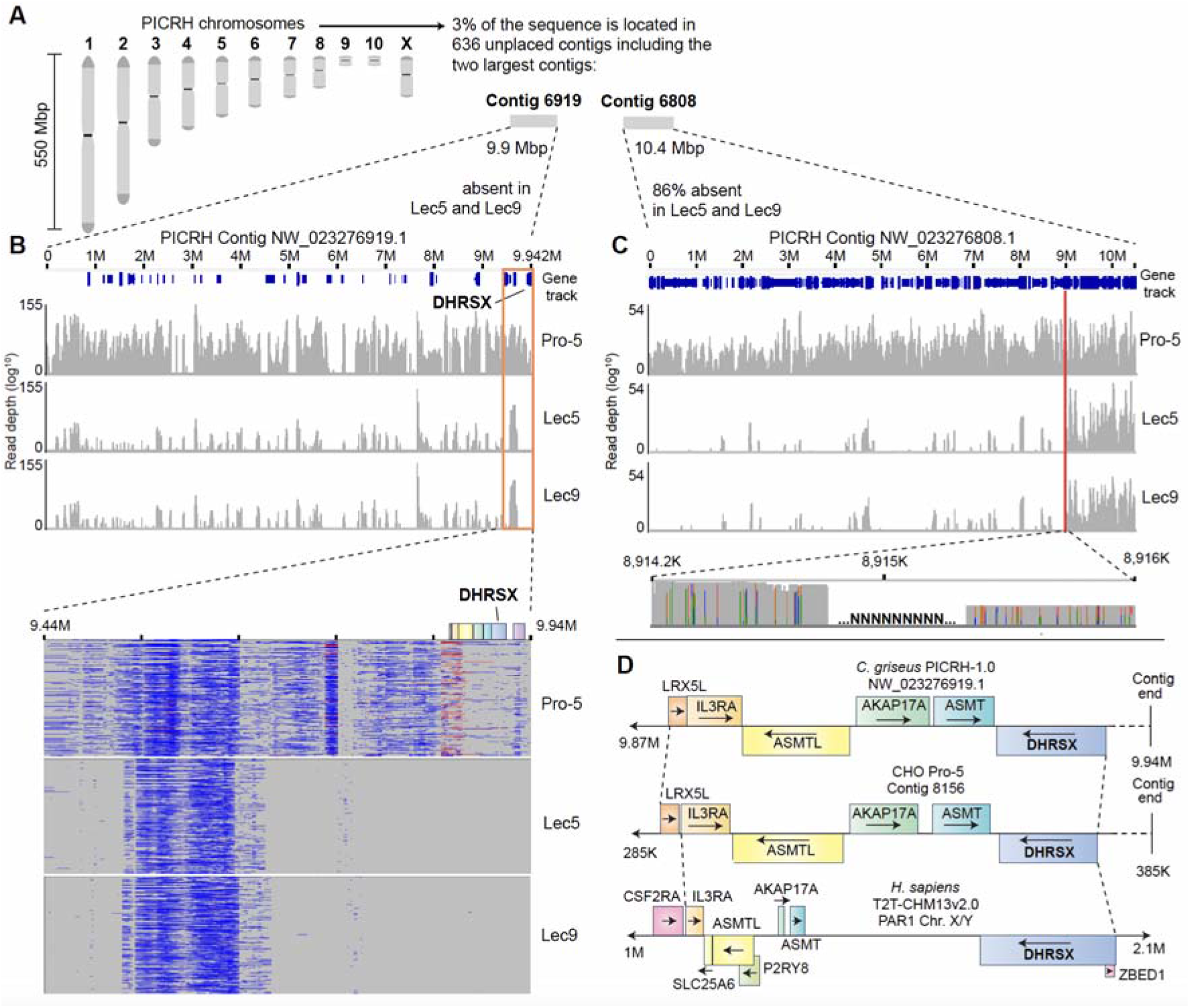
**The *DHRSX* gene is absent in Lec5 and Lec9 CHO cells** (A) The Chinese hamster chromosomes. (B) Upper panel: The contig containing *DHRSX* is devoid of consistent reads in Lec5 and Lec9 CHO cells, but present in parental Pro-5 cells. Chart shows the coverage (range 0-155, log^10^ scale) of reads recovered from ONT long-read whole genome sequencing of Pro-5, Lec5 and Lec9 CHO cells, mapped to contig NW_023276919.1 of the PICRH-1.0 (GenBank: GCA_003668045.2) *C. griseus* genome assembly. Coverage calculation (via igvtools count) and visualization performed using Integrative Genomics Viewer (IGV) 2.17.1 (46). Gene track = GCF_003668045.3. Lower panel: Reads from ONT sequencing of Pro-5, Lec5 and Lec9 CHO cells mapped to the terminal (9.44-9.94M) region of contig NW_023276919.1 of the PICRH-1.0 CHO genome. Visualization performed using Ribbon (47). (C) Upper Panel: The first 8,915 Kbp of the unplaced contig NW_023276808.1 are absent in Lec5 and Lec9 cells. Chart shows the coverage (range 0-54, log^10^ scale) of reads recovered from ONT long-read whole genome sequencing of Pro-5, Lec5 and Lec9 CHO cells, mapped to contig NW_023276808.1 of the PICRH-1.0 (GenBank: GCA_003668045.2) *C. griseus* genome assembly. Red line indicates position upstream of which reads are almost completely absent in Lec5 and Lec9 cells. Coverage calculation (via igvtools count) and visualization performed using Integrative Genomics Viewer (IGV) 2.17.1 (49). Gene track = GCF_003668045.3. Lower Panel: The gap of ±600bp containing N’s at approximately 8,915Kbp in Pro-5 reads mapped to PICRH NW_023276808.1 indicates that the reads up- and downstream of this location are not contiguous. Visualization performed using Integrative Genomics Viewer (IGV) 2.17.1. (D) Conservation of synteny in the genomic region surrounding *DHRSX* in the PICRH-1.0 *C. griseus* genome, contig 8156 derived from the *de novo* genomic assembly of Pro-5 CHO cells in this study, and *H. sapiens* T2T-CHM13v2.0. Note extended scale in *H. sapiens* (20x that of CHO genomes).

**Table 1:** Parameters of *de novo* assembly in this study compared to available *C. griseus* (PICRH) and CHO-K1 (CHOZN v2.3) reference genomes. N50: Length of the shortest contig at 50% of assembly length (i.e. 50% of assembly is in contigs of this length or longer); N90: Length of the shortest contig at 90% of assembly length (i.e. 90% of assembly is in contigs of this length or longer); L50: The minimum number of contigs that, combined, make up 50% of the assembly length.

| Assembly | This study | This study | This study | PICRH (2020) | CHOZN v2.3 (2022) |
| --- | --- | --- | --- | --- | --- |
| DNA Source | Wild type<br>Pro-5 CHO | Lec5 CHO | Lec9 CHO | *C. griseus*<br>tissues | CHO-K1 cells |
| Assembly length (Gb) | 2.39 | 2.35 | 2.35 | 2.37 | 2.30 |
| Contig N50 | 13,782,288 | 14,039,095 | 13,096,259 | 274,391,693 | 43,523,667 |
| Contig N90 | 1,667,153 | 1,803,015 | 1,796,527 | 127,255,434 | 549 |
| Contig L50 | 48 | 47 | 51 | 4 | 17 |
| No. of contigs | 3,013 | 2,119 | 2,241 | 647 | 1,634,314 |

Next, we mapped our ONT long-reads to the CriGri-PICRH-1.0 genome (subsequently referred to as PICRH). This is the Chinese hamster genome with the most complete chromosomal information, with 97% of the genome placed on 11 ‘mega-scaffolds’ pertaining to the Chinese hamster chromosomes (**Fig. 5A**) (27). Coverage of all 11 characterized chromosomes was normal in both Lec5 and Lec9 cells, when compared to Pro-5 cells (**Table S3**). The same was found when, instead of mapping ONT long-reads directly to the PICRH reference genome, we used the consensus reference sequences created by *de novo* assembly of each respective genome (**Table 2, Table S4**).

**Table 2:** Mean coverage of consensus sequence derived from *de novo* assembly of CHO Pro-5, Lec5 and Lec9 genomes when mapped to the chromosomes of the PICRH reference genome. See Table S4 for data from all PICRH contigs.

| PICRH Chromosome | Length (bp) | Pro-5 coverage (%) | Lec5 coverage (%) | Lec9 coverage (%) |
| --- | --- | --- | --- | --- |
| 1 | 550,089,852 | 100 | 100 | 100 |
| 2 | 461,620,117 | 100 | 100 | 100 |
| 3 | 282,827,514 | 100 | 100 | 100 |
| 4 | 231,097,868 | 100 | 100 | 100 |
| 5 | 193,770,019 | 099 | 099 | 099 |
| 6 | 155,611,870 | 100 | 099 | 099 |
| 7 | 134,359,064 | 100 | 100 | 099 |
| 8 | 99,554,469 | 113 | 111 | 111 |
| 9 | 28,505,831 | 103 | 107 | 102 |
| 10 | 32,558,357 | 101 | 101 | 100 |
| X | 127,255,434 | 103 | 102 | 103 |

In the PICRH genome, *DHRSX* is located close to the end of an unplaced contig of approximately 10 Mbp (NW_023276919.1). The mapping of Pro-5 reads to this contig resulted in consistent coverage. However, our long-read data sets from Lec5 and Lec9 cells left this contig empty, except for reads in repetitive regions that are unlikely to be representative of real coverage or alignment. This indicated that this region is entirely absent in the mutant cell lines (**Fig. 5B**), in turn suggesting that both the Lec5 and Lec9 cell carried a large deletion of at least 10 Mbp.

Several observations indicate that the deletion is in fact larger: First, approximately 86% of an additional unplaced contig of 10.5 Mbp, NW_023276808.1, was absent in Lec5 and Lec9 cells (**Fig 5C**). While the abrupt end of read alignments might give the impression of a distinct breakpoint in Lec5 and Lec9 cells, this transition occurs in a region that contains unidentified sequences (‘NNN’ in **Fig 5C**) and is not spanned by any reads, suggesting that the assembly of this contig is incorrect. Second, Lec5 and Lec9 cells lack reads in many other unplaced contigs (see **Tables S2 and S3**), suggesting that the deletion is likely even larger. Given that no major differences could be identified in the gross coverage of any contig between Lec5 and Lec9 cells, deletions in both cells were likely spanning a very similar region that exceeds 19 Mbp.

To attempt a better localization of *DHRSX* compared to a deletion breakpoint or telomere, we explored the synteny of this region between PICRH contig NW_023276919.1, contig 8156 in our *de novo* assembly of the Pro-5 cell genome, and corresponding human sequences. Indeed, the organization and directionality of gene transcripts in the terminal regions of these two contigs are conserved extremely well between hamster assemblies (**Fig. 5D**). The same is largely true in *H. sapiens*, though, intriguingly, the intergenic and intronic sequences were greatly expanded (∼20x) compared to those in PICRH and Pro-5 cells. Thus, in Lec5 and Lec9 cells, *DHRSX* and its genetic neighborhood (containing *LRX5L, IL3RA, ASMTL, AKAP17A,* and *ASMT*) is entirely absent **(Fig. 5B & 5D**).

The closest gene to *H. sapiens DHRSX* in the centromeric direction is *CD99* (alongside its pseudogene *CD99P1)*. *CD99* is found close to the start of another short PICRH contig of 120kb (NW_023277415.1), also missing in Lec5 and Lec9 cells. In our mapping, overlapping reads in Pro-5 cells between NW_023276919.1 and this contig indicate that they are also contiguous in *C. griseus*, but genes further along this contig do not correspond to their human orthologues (*XG, GYG2, ARSD, ARSE, ARSL*). Given the unplaced nature of the hamster contigs, we were not able to determine whether DHRSX was located close to the chromosomal end or close to the end of the deletion.

Finally, to confirm that chDHRSX was active and catalyzed the same activities as hDHRSX, we overexpressed and purified recombinant chDHRSX using the amino acid sequence predicted from whole genome long-read sequencing of Pro-5 CHO cells (**Fig. S2**). When assessing polyprenol dehydrogenase and dolichal reductase activities, we observed kinetic parameters comparable with those observed for hDHRSX (**Fig. 6**, **Table 3**).

**Figure 6.**
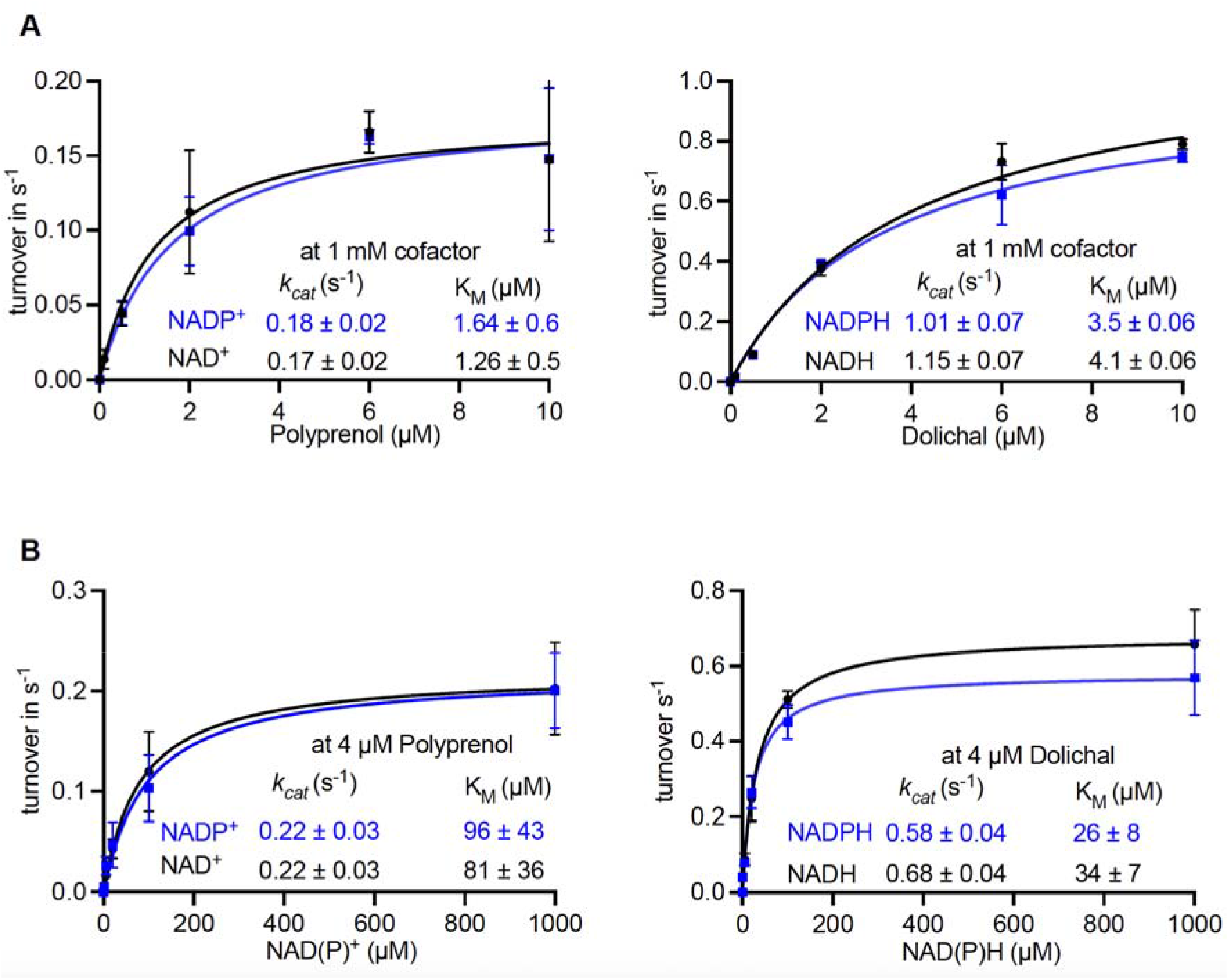
**Chinese hamster DHRSX is functional with similar kinetic parameters to the human ortholog** (A) For polyprenol and dolichal: Kinetic parameters were determined by measuring polyprenal or dolichol formation after incubation of the indicated concentrations of polyprenol or dolichal, respectively. Assays were performed in the presence of 1 mmol/L of the indicated cofactors and 0.01 μmol/L recombinant chDHRSX for 5 min at 37 °C. Data are turnover rates based on formation of polyprenal-18 or dolichol-18 (means ± SEM; n=3). (B) For NAD(P)^+^ and NAD(P)H: identical incubation was performed with variable nucleotide concentrations in the presence of 4 µmol/L polyprenol or dolichal and measuring the formation of polyprenal or dolichol, respectively. Data are turnover rates based on formation of polyprenal-18 or dolichol-18 (means ± SEM; n=3).

**Table 3:** Kinetic properties of chDHRSX determined in this study and those of hDHRSX previously reported (See ref. 18).

|  |  | Polyprenol | Dolichal | NAD<sup>+</sup> | NADP<sup>+</sup> | NADH | NADPH |
| --- | --- | --- | --- | --- | --- | --- | --- |
| ***K<sub>cat</sub>***<br>**(s<sup>-1</sup>)** | **chDHR SX**<br>*(This study)* | 0.17 ± 0.02 | 1.01 ± 0.07 | 0.22 ± 0.03 | 0.22 ± 0.03 | 0.68 ± 0.04 | 0.58 ± 0.04 |
|  | **hDHR SX**<br>*(See ref. 18)* | 0.44 ± 0.05 | 1.00 ± 0.13 | 0.16 ± 0.03 | 0.18 ± 0.01 | 0.62 ± 0.08 | 0.65 ± 0.05 |
| ***K<sub>M</sub>***<br>**(mM)** | **chDHR SX**<br>*(This study)* | 1.3 ± 0.5 | 3.5 ± 0.1 | 81 ± 36 | 96 ± 43 | 34 ± 07 | 26 ± 08 |
|  | **hDHR SX**<br>*(See ref. 18)* | 5.7 ± 1.2 | 2.2 ± 1.3 | 66 ± 43 | 42 ± 11 | 45 ± 24 | 40 ± 12 |

## Discussion

### Solving an old riddle

Lec5 and Lec9 cells are characterized by a deficient conversion of polyprenol to dolichol (3–8) but the genetic basis of this phenotype was not identified. With the discovery of the role of SRD5A3 in the conversion of polyprenol to dolichol (17), a likely candidate for the molecular defect in Lec5 and Lec9 cells was revealed. However, it took the discovery of the role of DHRSX in the formation of dolichol (18), alongside the reassignment of the function of SRD5A3, to allow us to identify, here, the biochemical and genetic bases of the dolichol defect in Lec5 and Lec9 cells.

### A comparable large deletion comprising *DHRSX* causes the glycosylation defect in Lec5 and Lec9 cells

Lec5 and Lec9 cells have been postulated to have the same gene defect since Lec5/Lec9 hybrid cells produce polyprenol rather than dolichol (16, 28). However, earlier experiments showed Lec5/Lec9 hybrids are sensitive to ricin, indicative of complementation and a different genetic basis for each phenotype (29). A compensatory mechanism that allows ricin resistance in hybrids might exist, via a similar mechanism to that observed in certain cells from DHRSX-CDG patients (18). Alternatively, ricin resistance in CHO cells can arise from mutations affecting protein synthesis (30) which may be present in either Lec5 or Lec9 cells.

Nevertheless, we show here that both Lec5 and Lec9 have a large deletion of at least 19 Mbp, leading to a loss of *DHRSX*, alongside many other genes, in contigs NW_023276919.1 and NW_023276808.1 of the PICRH *C. griseus* genome assembly. It appears therefore that the *DHRSX* gene is prone to inactivation by dramatic rearrangements, consistent with the fact that the *DHRSX* RNA was not identified in transcriptomics studies on Lec5 cells by Lu *et al.* (31). Given that both cell lines have a comparable deletion, a similar deletion may also have occurred in the several other cell lines that spontaneously arose and that fall in the same complementation group as Lec5 or Lec9 cells (e.g. CHB11-1-3 cells (16)). All these cell lines were selected for their resistance to toxic lectins that bind to N-glycans. Thus, even if this deletion might occur at an extremely low rate (the genomic instability of CHO cells is well documented (32, 33)) and is unfavorable to growth, it likely confers a selective advantage under the significant selective pressure applied by the lectin.

### CHO Lec5 and Lec9 revertants: an ongoing puzzle

The glycosylation phenotype of Lec5 and Lec9 cells is quite puzzling, since spontaneous revertants have been repeatedly observed (1, 29, 31). The large genomic deletion observed in these cell lines excludes the possibility that DHRSX might be reactivated. To account for this easy reversibility, Lu *et al.* suggested that the mutation(s) causing the Lec5 phenotype were affecting the epigenetic regulation of one or more genes; specifically, via homeobox genes. Our findings do not exclude this possibility. Conceivably, epigenetic modulation of one of the multiple dehydrogenases belonging to the same family as DHRSX (e.g. RDH11-14, DHRS7, DHRS13, WWOX), could lead to sufficient polyprenol dehydrogenase and dolichal reductase activity to restore sufficient glycosylation. The Lec5 revertant RNA-Seq data collected by Lu *et al.* does not show increased expression of any of these genes specifically. Another possibility for reversion might be the occurrence of mutations altering the kinetic properties in one of the many dehydrogenases belonging to the same family as DHRSX and retinol-dehydrogenases.

Alternatively, the fact that revertants of CHB11-1-3 (Lec5) cells synthesize normal levels of dolichol, but with higher levels of polyprenol (16) suggests that perhaps the polyprenol biosynthesis pathway (derived from the mevalonate/cholesterol biosynthesis pathway) is somehow upregulated in revertants. This might increase polyprenol levels to those high enough that sufficient N-glycosylation is restored, thereby diminishing their lectin-resistant phenotype. However, the opposite of this was identified in the gene expression data of Lu *et al.*, with mevalonate pathway genes such as 3-hydroxy-3-methylglutaryl-CoA synthase 1; (*Hmgcs1)* downregulated in Lec5 revertants.

It is likely that if another dehydrogenase can replace chDHRSX, it has a higher K_m_ for polyprenol. This situation resembles that in human cells: loss of DHRSX or SRD5A3 in both model cell lines and patient cells such as fibroblast and lymphoblasts does not result in the complete absence of dolichol. In some cases, levels are almost completely normalized, despite persistent polyprenol or polyprenal accumulation (18, 34). This suggests that the defects in either of these enzymes can be compensated at the cellular level by a yet unknown mechanism. Perhaps the compensatory mechanism in Lec5 and Lec9 revertants is, in fact, replicated in human cells, but not enough to prevent disease.

To what extent this type of compensation mechanism occurs widely in different tissues, and can attenuate specific symptoms, would certainly be worth investigating. Of note, the defect in the glycosylation of transferrin present in both DHRSX-CDG or SRD5A3-CDG can be normal or normalize with age (18, 35). Understanding the mechanism of reversion of the defect in CHO cells could also apply to human disease.

## Experimental procedures

### CHO cell culture

CHO cells were cultured in DMEM/F12 (Gibco, 31330-038) supplemented with 10% FBS (Clone III, HyClones) and antibiotics streptomycin (100 μg/mL), penicillin (100 U/mL), and amphotericin (0.5 μg/mL) at 37°C under 5% CO_2_. Cells were passaged using trypsin (Gibco, 25300096) and versene (Gibco, 15040066).

### Plasmid Constructions

The recombinant plasmid pET15b chDHRSX-N-His, pET15b hDHRSX-N-His (NM_145177.3) and pcDNA3.1(+) hSRD5A3-N-His (NM_024592.5) were purchased from GenScript Biotech (Rijswijk, Netherlands) and sequences were checked by Sanger sequencing. The primer sets hDHRSX_sense_BglII_HIS/hDHRSX_rev_BsrGI or SRD5A3_sense_BglII_HIS/SRD5A3_rev_bsrGI, respectively were used to PCR-amplify hDHRSX or hSRD5A3 from pET15b hDHRSX-N-His and pcDNA3.1(+) hSRD5A3-N-His using Phusion™ DNA polymerase, 2 U/μL (F-530S, ThermoFisher scientific) following the recommendations of the manufacturer. BgIII and BsrGI restriction enzymes were used for cloning into pUB83. T4 DNA ligase, 5 U/μL (EL0014, ThermoFisher scientific) was used for the insertion of hDHRSX or hSRD5A3 in pUB83 predigested by the same restriction enzymes. pUB83 is a lentiviral expression vector based on the pLVX-PURO (Clontech) plasmid containing a CMV promoter. It contains ampicillin and puromycin resistance cassettes for bacterial and eukaryotic cell selection, respectively. Sequence of recombinant vector pUB83 hDHRSX-N-His and pUB83 hSRD5A3-N-His were verified by Sanger sequencing. The pcDNA3-ER-Halo3N construct has been described previously (21).

### Recombinant lentiviruses production and infection of CHO cells

Recombinant lentiviruses were produced in HEK293T cells as previously described (36)using recombinant vector pUB83 hDHRSX-N-His, pUB83 hSRD5A3-N-His or empty pUB83. Viruses were then used to infect Pro-5, Lec5 or Lec9 cells. CHO cells were grown in 6 well plates to reach 60 % confluence on the day of infection in DMEM/F12. The medium was replaced by DMEM/F12 containing diluted recombinant lentiviruses (lentivirus: DMEM 1:4 V/V) in the presence of 4 µg/mL polybrene. After 24 hours of growth the medium was replaced by virus free DMEM/F12 containing 30 µg/mL puromycin.

### Isoprenoid species and preparation of dolichal

Dolichol (# 9002000), polyprenol (# 9002100) and polyprenal (# 9002200) were purchased from Avanti Polar Lipids. Dolichal was synthesized by oxidizing dolichol in the presence of pyridinium dichromate(37) (# 214698-100G, Merck life science BV): 300 mL of 1 mg/mL dolichol in chloroform was dried down and resuspended in 200 mL of dichloromethane containing 10 mg/mL of pyridinium dichromate. After shaking at 1000 rpm for 15 min, the preparation was incubated overnight at 20 °C. The mixture was centrifuged, and the supernatant was transferred to a new tube. Several cycles of back-extractions were performed by using 200 mL methanol/water (5:3) (MS-grade, Biosolve) to remove pyridinium dichromate until the preparation became translucent. The quality of dolichal synthesis was assessed by LC-MS, showing a high yield of conversion (> 95 %). Dolichal was stored at -80 °C for future enzymatic assays. All isoprenoid solutions represent mixtures with 13 to 21 isoprene units (with 18 being the most abundant species). Molar concentrations were calculated based on the distribution of chain lengths. Thus, a 5 µg/mL solution corresponds to approximately 4 µM.

### Sample preparation and extraction of metabolites from CHO cells

The medium was removed, plates were rapidly washed with ice-cold water and plunged in liquid nitrogen to quench metabolic activity. The frozen dishes were placed on dry ice, and 500 µL ice-cold methanol was immediately added, followed by 300 µL of ice-cold water. The cells were scraped and collected in 2 ml tubes containing 1 ml of chloroform (MS-grade, Biosolve). Tubes were vigorously mixed at 2000 rpm for 20 min at 4°C, followed by a centrifugation at 13200 rpm for 30 min at 4°C. The lower layer, containing hydrophobic metabolites, was dried down under a gentle stream of nitrogen and resuspended in 100 µL of methanol/isopropanol (1:1) before LC–MS analysis.

### Dimethylation of isoprenoid phosphates using trimethylsilyl diazomethane (TMSD)

The dimethylation of dolichol phosphate and polyprenol phosphate was performed as described in Kale et al., (2023) (38): The organic fraction containing isoprenoid species was completely dried and resuspended in 200 μL of dichloromethane:methanol (6.5:5.2, v/v). 10 μL of 2 mol/L TMSD was added and incubated for 40 min at room temperature. The incubation was halted by adding 1 μL of glacial acetic acid. Samples were then dried and dissolved in methanol:isopropanol 1:1 (V/V) for LC-MS analysis.

### Extraction of membrane proteins from CHO cells

Confluent 10 cm plates were washed with ice-cold PBS and cells were collected in 0.5 ml/plate of lysis buffer (25 mmol/L HEPES pH 7.5, 2 μg/mL antipain and leupeptin, 0.5 mmol/L PMSF and 10% glycerol) by scraping plates. Cells were then subjected to 2 cycles of freezing-thawing in liquid nitrogen, treated with DNAse I (0.1 mg/mL in 10 mmol/L MgSO_4_) for 15 min and centrifuged (13200 rpm for 15 min at 4°C) to recover a membrane fraction in the pellet. The resulting pellet was washed twice by centrifugation as above with the same volume of lysis buffer and the resulting pellet was resuspended in the initial volume of lysis buffer. This preparation was aliquoted and kept at -80°C.

### Measurement of DHRSX or SRD5A3 activities in membrane protein extracts of CHO cells

Enzymatic activity measurements were performed in buffer containing 10 mmol/L HEPES, pH 8.0, 10 mmol/L KCl, 0.3 % Triton X-100, 0.5 mmol/L β-mercaptoethanol, 1% phosphatidylcholine (PC), and 0.2% phosphatidylethanolamine (PE). To assess dehydrogenase or reductase activities, 5 μg/mL of polyprenol, polyprenal or dolichal were used in the presence of 5 mmol/L of NAD(H)^+^ or NADP(H)^+^. First, 5 μL of a of 50 µg/mL solution of the respective polyisoprenoid mixtures in chloroform was added into an empty tube and dried under a stream of nitrogen. After that, 5 μL of 10X buffer [100 mmol/L HEPES, pH 8.0, 100 mmol/L KCl, 3% Triton X-100, 0.5 mmol/L β-mercaptoethanol, 10% PC, and 2% PE] were added to the tube. 5 uL of NAD(P)^+^ or NAD(P)H (50 mM) was then added. The incubations were started by adding protein extracts at a final concentration of 1 mg/mL in a total volume of 50 µL. Incubation was performed at 37 °C for 2h. Samples were analyzed by LC-MS after methanol-chloroform extraction of analytes.

### Analysis of recombinant chDHRSX activity on polyprenol and dolichal

To determine kinetic properties of recombinant chDHRSX, measurements were performed in a buffer containing 10 mmol/L HEPES pH 8.0, 10 mmol/L KCl, 0.5 mmol/L β-mercaptoethanol, 1% PC, and 0.2% PE. 4 μmol/L of lipid substrates were used in the presence of 0, 1, 5, 20, 100, or 1000 μmol/L of NAD(P)^++^ or NAD(P)H to investigate kinetic properties of cofactors. For the ones of polyprenol and dolichal: 0, 0.1, 0.5, 2, 6 or 10 μmol/L of lipids were used in the presence of 1 mmol/L of NAD(H). After a preheating at 37°C, the assay was started by addition of 0.01 μmol/L final concentration of recombinant chDHRSX and carried out in a total volume of 50 μL at 37°C and 400 rpm. After five minutes of incubation, assays were immediately quenched by methanol-chloroform extraction. Non-linear curve fitting with GraphPad Prism 10.1 was used assuming Michaelis Menten kinetics.

### Sample preparation and extraction of metabolites from enzymatic assays

After the end of each respective incubation, 450 µL of ice-cold methanol/water (5:3) was added into the tube followed by 550 µL of ice-cold chloroform. The mixture was vigorously mixed at 2000 rpm for 20 min and at 4°C, followed by a centrifugation at 13200 rpm for 10 min at 4°C. The lower layer containing hydrophobic metabolites was dried down under a gentle stream of nitrogen and resuspended in 50 µL methanol/isopropanol (1:1) before LC–MS analysis.

### Stable expression of pcDNA3-Halo3N-ER

Pro-5, Lec5 or Lec9 CHO cells already overexpressing empty pUB83 or hDHRSX pUB83 were reverse transfected using the Lipofectamine 3000 reagent (Thermo Fisher Scientific) with 0.5 μg of the pcDNA3-Halo3N-ER plasmid into 6-well plates containing DMEM/F-12 and 30 µg/mL puromycin. After 24 hours, medium was swapped with that also containing 1 mg/mL G-418 (Sigma, A1720) for selection. For treatment with Tunicamycin (Sigma, SML1287) and MG-132 (VWR, 474787), cells were plated on 6-well plates and after 24 hours medium was exchanged for that also containing 1μg/mL Tunicamycin and/or 10 μmol/L MG-132. Cells were cultured for a further 24 hours before collection of lysates for immunoblot analysis.

### SDS-PAGE and western blot

For immunoblotting, protein lysates from CHO cells were prepared by lysing cells in RIPA buffer (10 mmol/L Tris-HCl [pH 7.4], 150 mmol/L NaCl, 0.5% sodium deoxycholate, 0.1% SDS, and 1X cOmplete protease inhibitor cocktail [Sigma Aldrich]) at 4°C, passing 5 times through a 27G needle, incubating for 30 min followed by centrifugation at 4°C (15,000 g, 30 min). Protein concentration was determined with the Pierce BCA protein assay kit (Thermo Fisher Scientific, 23225). Lysates were separated by SDS-PAGE and blotted onto a nitrocellulose membrane (Invitrogen, LC2000). Blocking was performed in 2-5% milk or bovine serum albumin (98022, Sigma-aldrich) with the relevant primary and secondary antibodies. Washing was performed with 1x Tris-buffered saline solution (12498S, Bioké) with 0.1% Tween-20 (P1379, Sigma-Aldrich). Signal detection was performed by chemiluminescence using an Amersham ImageQuant 800 imager (Cytiva), and quantification using ImageJ 2.14.0. Primary antibodies used were anti-DHRSX (Sigma, HPA003035), anti-β-tubulin (Thermo Fisher Scientific, MA516308), anti-HaloTag® (Promega, G9211) and secondary antibodies were HRP-linked anti-rabbit (7074S) or mouse (7076S) IgG (Bioké).

### LC–MS analysis of metabolites

LC–MS analysis of organic fractions obtained from cells (or enzymatic assays) was carried out using a method adapted from that used by Dewulf *et al.* 2019 (39). Briefly, 5 µL of sample was injected and subjected to reverse phase chromatography with an Accucore C30 150 x 2.1 mm column (ref 27826-152130, Thermo Fisher Scientific), operated at 45 °C on an Agilent 1290 HPLC system. The flow rate was constant at 0.2 mL/min using mobile phase A (60% acetonitrile, and 40% water, 10 mmol/L ammonium formate, and 0.1% formic acid) and B (90% isopropanol, 10% acetonitrile, 10 mmol/L ammonium formate, and 0.1% formic acid (Biosolve)). An Agilent 6546 ion funnel mass spectrometer was used in the positive or negative ionization modes with an electrospray ionization (ESI) (voltage 3500 V, Nozzle voltage 1000 V, sheath gas 350°C at 11 L/min, nebulizer pressure 35 psi and drying gas 300°C at 8 L/min). Starting from 5 min onwards, one spectrum encompassing a range of 69 to 1700 *m/z* was acquired per second, generated from 10772 transients. The mass spectrometer was operated in positive polarity for the detection of dolichal, polyprenal, dolichol, polyprenol, dimethylated dolichol-P, and dimethylated polyprenol-P. For the elution, the solvent gradient was: 0–5 min at 90% B; 5–33 min from 90 to 97% B; 33–34 min from 97 to 99% B; 34–35 min from 99 to 90% B. The negative polarity was used to detect and measure polyprenoic acid, dolichol-P, polyprenol-P, dolichol-P-hexose, and polyprenol-P-hexose. The elution gradient consisted of: 0–3 min at 30% B; 3–8 min from 30 to 43% B; 8–9 min from 43 to 50% B; 9–18 min from 50 to 90% B; 18–26 min from 90 to 99% B; 26–30 min at 99% B; 30–30.1 min from 99 to 30% B; 30.1–35 min at 30% B. The different theoretical *m/z* values of [M + NH_4_^+^] and [M - H^+^] ions are given in **Table S5**. The resulting data were analyzed and processed by the software Agilent MassHunter Qualitative Analysis 10.0 for the identification and the visualization of peaks/metabolites. The quantification was performed by Agilent MassHunter Quantitative Analysis software (Agilent Technologies, CA, USA).

### Proteomic analysis

40 µg of membrane preparations were resuspended in 46 µL of a solution containing 5% SDS, 50 mM triethylammonium bicarbonate buffer pH 8.5 (TEAB) and 10 mM tris(2-carboxyethyl)phosphine (TCEP). After 10 min at 95°C, chloroacetamide was added to a final concentration of 20 mM followed by another 30 min at RT. Samples where then acidified by addition of phosphoric acid to a final concentration of 2.5%, split into 2 parts and transferred onto a micro S-Trap column (Protifi LLC, USA). Digestion was performed at 37°C overnight using 1:100 of Trypsin and LysC, either with or without 1 µL of PNGase F (500 Units, NEB).

Digested peptides were eluted in three steps using 40 µL each of 50 mM Tris pH 8.5, 0.2% formic acid and 50% acetonitrile. The eluted peptides were dried down in a vacuum concentrator (SpeedVac, ThermoFisher Scientific) and resuspended in 2% acetonitrile and 0.2% formic acid. Peptide concentration was determined by Pierce™ Quantitative Peptide Assay (ThermoFisher Scientific). Peptides were directly loaded onto reversed-phase trap-column (Acclaim PepMap 100, ThermoFisher Scientific) and eluted in backflush mode. Peptide separation was performed using a reversed-phase analytical column (EasySpray, 0.075 x 250 mm, ThermoFisher Scientific) with a linear gradient of 4%-27.5% solvent B (0.1% formic acid in 98% acetonitrile) with solvent A (0.1% formic acid) for 100 min, 27.5%-40% solvent B for 10 min, 40%-95% solvent B for 10 min at a constant flow rate of 300 nL/min on a Vanquish Neo HPLC system. The peptides were analyzed by an Orbitrap Fusion Lumos tribrid mass spectrometer with an ESI source (ThermoFisher Scientific) coupled online to the nano-LC and operated in positive polarity. Peptides were detected in the Orbitrap at a resolution of 120,000. Peptides were selected for MS/MS using the HCD setting at 30; ion fragments were detected in the Orbitrap at a resolution of 30 000. A data-dependent procedure was used with a cycle time of 3 s alternating between one MS scan and multiple MS/MS scans for ions above a threshold ion count of 38000 in the MS survey scan with 30 s dynamic exclusion. The electrospray voltage applied was 2.1 kV. MS1 spectra were obtained with an AGC target of 400000, and a maximum injection time of 100 ms. MS2 spectra were acquired with an AGC target of 100000 and an automatic adjustment of the maximum injection time. ms. The *m/z* scan range was 350 to 1800 for MS, and automatically adjusted for MS/MS scans according to the precursor ion mass.

For samples without PNGase F treatment, a modified version with a second MS/MS step was used. When specific masses were observed in the first MS/MS step, EThcD fragmentation was triggered to facilitate the localization of the glycosylation site. This included m/z 204.0867 for HexNac, m/z 138.0545 for a HexNAc fragment, m/z 366.1396 for HexNac-Hex, m/z 292.102 for sialic acid, m/z 163.06 for Hex, and m/z 454.16 for HexNeuAc. The supplemental activation energy in EThcD was set to 30%, and ion fragments were detected in the Orbitrap at a resolution of 30 000, with an AGC target of 300 000, a maximum injection time of 250 ms, and an *m/z* scan range from 150 to 2000.

The resulting MS/MS data were processed using the Sequest HT search engine within Proteome Discoverer 2.5 SP1 against a *Cricetulus griseus* protein database obtained from Uniprot (sp_canonical TaxID=10029) with the addition of the human DHRSX sequence. Trypsin was specified as cleavage enzyme allowing up to 2 missed cleavages, 4 modifications per peptide and up to 7 charges. The mass error was set to 20 ppm for precursor ions and 0.05 Da for fragment ions. Oxidation on Met (+15.995 Da), Hex(1)HexNAc(2) (+568.212 Da), Hex(2)HexNAc(2) (+730.264 Da), Hex(3)HexNAc(2) (+892.317 Da), Hex(4)HexNAc(2) (+1054.370 Da), Hex(5)HexNAc(2) (+1216.423 Da), Hex(6)HexNAc(2) (+1378.476 Da), Hex(7)HexNAc(2) (+1540.529 Da), Hex(8)HexNAc(2) (+1702.581 Da), Hex(9)HexNAc(2) (+1864.634 Da), Hex(10)HexNAc(2) (+2026.687 Da), Asn->Asp (+0.984 Da) on Asparagine, and Methionine loss (-131.040 Da) on the N-terminus of the protein and peptides were considered as variable modifications accordingly, whereas carbamidomethylation of cysteine was considered as a fixed modification. The False discovery rate (FDR) was assessed using Percolator and thresholds for the identification of proteins, peptides and modification sites were specified at 1%. Label-free quantification of peptides is based on the precursor ion intensity. Signals were normalized to the sum of all signals within each individual sample. Protein abundances were calculated as the sum of the abundances of unmodified peptides. To facilitate visualization in a heat map in Graphpad Prism, data were normalized to the 70^th^ percentile of the abundance of the same peptide within the different samples.

### Long-read sequencing and assembly

High molecular weight genomic DNA from Pro-5, Lec5 and Lec9 CHO cells was extracted using the Monarch^®^ HMW DNA Extraction Kit for Cells & Blood (NEB, T3050L). Resulting gDNA was then sheared by passing each sample 25 times through a 26G needle. Average fragment lengths of 40-50 kbp were confirmed using a Fragment Analyzer (Advanced Analytical) with an HS Large Fragment 50 Kb analysis kit (Agilent, DNF-464-33). Samples were prepared for Oxford Nanopore Technologies (ONT) long-read sequencing using a Ligation Sequencing Kit XL V14 (ONT, SQK-LSK114-XL) before loading of 1μg DNA onto a R10.4.1 PromethION Flow Cell (ONT, FLO-PRO114M). Run quality control was performed using ONT’s MinKNOW Core 5.4.3. Resulting raw data was base called using Dorado 0.1.1 with the super accurate model. Raw statistics were obtained using NanoPlot 1.41.0 (40). Assembly was performed using flye 2.9.2 (41) and polished with racon 1.5.0 (42). Assembly quality was assessed with QUAST 5.2.0 (43). For mapping of flye-derived *de novo* assembly contigs or consensus sequences to the PICRH genome, Minimap 2.2.25 (44) was used for mapping of ONT reads and flye-derived *de novo* assembly contigs to the PICRH genome, with the models map-ont and asm20, respectively. Coverage was computed using Mosdepth 0.3.5 (45) and compiled using RStudio v4.3.2. Coverage and assembly data was analyzed and/or visualized using Integrative Genomics Viewer (IGV) 2.17.1 (46), Ribbon (47)and NCBI’s Genome Workbench v4.9.1.

### Overexpression and Ni-NTA agarose purification of chDHRSX in *E. coli*

pET15b plasmids containing the cDNA sequence of *chDHRSX* with a 6 x His tag appended to the N-terminus were initially transformed into XL1 blue chemically competent *E. coli* (Life technologies, ref: C404003). Clones were selected and grown to prepare minipreps. Vectors were then transformed into BL21(DE3) *E. coli.* colonies were selected according to ampicillin resistance and grown in 10 mL LB medium (200 μg/mL ampicillin) overnight at 37°C and 200 rpm before dilution in 200 mL LB medium (200μg/mL ampicillin) and further growth until an OD_600_ of 0.5 was reached. Isopropylthio-β-galactoside (IPTG) was then added to a final concentration of 1 mmol/L and the culture was grown for a further 20 hours at 20°C and 200 rpm before centrifugation at 4000 x g for 20 minutes to harvest cells. Native purification of 6 x His-tagged chDHRSX was performed using the AKTA purifier 900 series (GE Healthcare) by using a Ni^2+^-resin column (HisTrap-HP 1 ml, GE Healthcare) as described previously (48).

Protein concentration in the purified preparations was estimated by measuring A_280_ and computing the concentration from the expected extinction coefficient (ε = 29450 M^-1^ cm^-1^) on the basis of the amino acid sequence (Protparam tool, at https://web.expasy.org/protparam/).

### Statistical analysis

All analyses were carried out via GraphPad Prism version 10.1 for Mac (GraphPad Software, La Jolla, CA). Statistical analysis was performed by a student’s T-test or one-way ANOVA followed by Tukey’s multiple comparisons test.

## Data availability

Mass spectrometry proteomics data have been deposited to the ProteomeXchange Consortium via the PRIDE partner repository with the dataset identifiers PXD052706 and 10.6019/PXD052706. ONT long-read sequencing data has been deposited to the NCBI Sequence Read Archive under the BioProject accession number PRJNA1120760.

## Supporting information

This article contains supporting information.

## Supporting information

Key resources table

## Acknowledgments

Thanks to Greet Peeters of the Laboratory for Cytogenetics and Genome Research and Wim Meert from UZ Leuven Genomics Core for advice on long-read sequencing whole genome analysis, as well as Didier Vertommen for advice on glycoproteomic analyses.

## Funding and additional information

The French National Agency (ENIGMncA project, ANR-21-CE14-0049-01) to F.F.; the CNRS IRP GLYCOCDG project to F.F; an FWO senior postdoctoral fellowship (Project ID: 1289023N to M.P.W.); the Jaeken-Theunissen CDG Fund; Mizutani Foundation for Glycoscience (Grant 210119 to G.M. and 240097 to G.T.B.); CELSA fund grant (CELSA/21/027) to G.M.), KU Leuven Global PhD Partnership (to C.R.A., F.F and G.M. 3M200250). WELBIO 2019 (to G.T.B.), ERC consolidator grant #771704 (to G.T.B.), Fondation Médicale Reine Elisabeth (to G.T.B.), FNRS equipment grant and research credit (to G.T.B.), ARC UCLouvain (to G.T.B.), FWO-FNRS WEAVE program (G061524N to G.T.B., M.P.W and G.M.), Fonds Baillet-Latour (to G.T.B. and E.V. S.) and an NIH grant R01 GM105399 to P.S.

## Conflict of interest

The authors declare that they have no conflicts of interest with the contents of this article.

## Notes

https://proteomecentral.proteomexchange.org/cgi/GetDataset?ID=PXD052706

http://www.ncbi.nlm.nih.gov/bioproject/1120760

### Competing Interest Statement

The authors have declared no competing interest.

## References

1. Patnaik, S. K., and Stanley, P. (2006) Lectin-resistant CHO glycosylation mutants. Methods Enzymol. 416, 159–182

2. Esko, J. D., and Stanley, P. (2015) Glycosylation Mutants of Cultured Mammalian Cells. Essentials of Glycobiology. 10.1101/GLYCOBIOLOGY.3E.049

3. Krags, S. S., Robbins, P. W., and Baker, R. M. (1977) Reduced Synthesis of [14ClMannosyl Oligosaccharide-lipid by Membranes Prepared from Concanavalin A-resistant Chinese Hamster Ovary Cells*. Journal of Biological Chemistry. 252, 3561–3564

4. Cifone, M. A., Hynes, R. O., and Baker, R. M. (1979) Characteristics of concanavalin A-resistant Chinese hamster ovary cells and certain revertants. J Cell Physiol. 100, 39–54

5. Rosenwald, A. G., Stanley, P., and Krag, S. S. (1989) Control of carbohydrate processing: increased beta-1,6 branching in N-linked carbohydrates of Lec9 CHO mutants appears to arise from a defect in oligosaccharide-dolichol biosynthesis. Mol Cell Biol. 9, 914–924

6. Rosenwald, A. G., and Krag, S. S. (1990) Lec9 CHO glycosylation mutants are defective in the synthesis of dolichol. J Lipid Res. 31, 523–533

7. Beck, P. J., Gething, M. J., Sambrook, J., and Lehrman, M. A. (1990) Complementing mutant alleles define three loci involved in mannosylation of Man5-GlcNAc2-P-P-dolichol in Chinese hamster ovary cells. Somat Cell Mol Genet. 16, 539–548

8. Krag, S. S. (1979) A Concanavalin A-resistant Chinese Hamster Ovary Cell Line Is Deficient in the Synthesis of [3H]Glucosyl Oligosaccharide-Lipid*. Journal of Biological Chemistry. 254, 9167–9177

9. Burda, P., and Aebi, M. (1999) The dolichol pathway of N-linked glycosylation. Biochim Biophys Acta. 1426, 239–257

10. Kean, E. L., Rush, J. S., and Waechter, C. J. (1994) Activation of GlcNAc-P-P-dolichol synthesis by mannosylphosphoryldolichol is stereospecific and requires a saturated alpha-isoprene unit. Biochemistry. 33, 10508–10512

11. Mclachlan, K. R., and Krag, S. S. (1992) Substrate specificity of N-acetylglucosamine 1-phosphate transferase activity in Chinese hamster ovary cells. Glycobiology. 2, 313–319

12. D’Souza-Schorey, C., McLachlan, K. R., Krag, S. S., and Elbein, A. D. (1994) Mammalian glycosyltransferases prefer glycosyl phosphoryl dolichols rather than glycosyl phosphoryl polyprenols as substrates for oligosaccharyl synthesis. Arch Biochem Biophys. 308, 497–503

13. McLachlan, K. R., and Krag, S. S. (1994) Three enzymes involved in oligosaccharide-lipid assembly in Chinese hamster ovary cells differ in lipid substrate preference. J Lipid Res. 35, 1861–8

14. Palamarczyk, G., Lehle, L., Mankowski, T., Chojnacki, T., and Tanner, W. (1980) Specificity of solubilized yeast glycosyl transferases for polyprenyl derivatives. Eur J Biochem. 105, 517–523

15. Stoll, J., Rosenwald, A. G., and Krag, S. S. (1988) A Chinese hamster ovary cell mutant F2A8 utilizes polyprenol rather than dolichol for its lipid-dependent asparagine-linked glycosylation reactions. Journal of Biological Chemistry. 263, 10774–10782

16. Quellhorst, G. J., Hall, C. W., Robbins, A. R., and Krag, S. S. (1997) Synthesis of dolichol in a polyprenol reductase mutant is restored by elevation of cis-prenyl transferase activity. Arch Biochem Biophys. 343, 19–26

17. Cantagrel, V., Lefeber, D. J., Ng, B. G., Guan, Z., Silhavy, J. L., Bielas, S. L., Lehle, L., Hombauer, H., Adamowicz, M., Swiezewska, E., De Brouwer, A. P., Blümel, P., Sykut-Cegielska, J., Houliston, S., Swistun, D., Ali, B. R., Dobyns, W. B., Babovic-Vuksanovic, D., van Bokhoven, H., Wevers, R. A., Raetz, C. R. H., Freeze, H. H., Morava, É., Al-Gazali, L., and Gleeson, J. G. (2010) SRD5A3 Is Required for Converting Polyprenol to Dolichol and Is Mutated in a Congenital Glycosylation Disorder. Cell. 142, 203–217

18. Wilson, M. P., Kentache, T., Althoff, C. R., Schulz, C., Bettignies, G. de, Cabrera, G. M., Cimbalistiene, L., Burnyte, B., Yoon, G., Costain, G., Vuillaumier-Barrot, S., Cheillan, D., Rymen, D., Rychtarova, L., Hansikova, H., Bury, M., Dewulf, J. P., Caligiore, F., Jaeken, J., Cantagrel, V., Schaftingen, E. Van, Matthijs, G., Foulquier, F., and Bommer, G. T. (2024) A pseudoautosomal glycosylation disorder prompts the revision of dolichol biosynthesis. Cell. 0, 1–17

19. Hall, C. W., McLachlan, K. R., Krag, S. S., and Robbins, A. R. (1997) Reduced utilization of Man5GlcNAc2-P-P-lipid in a Lec9 mutant of Chinese hamster ovary cells: analysis of the steps in oligosaccharide-lipid assembly . J Cell Biochem. 67, 201–215

20. Los, G. V., Encell, L. P., McDougall, M. G., Hartzell, D. D., Karassina, N., Zimprich, C., Wood, M. G., Learish, R., Ohana, R. F., Urh, M., Simpson, D., Mendez, J., Zimmerman, K., Otto, P., Vidugiris, G., Zhu, J., Darzins, A., Klaubert, D. H., Bulleit, R. F., and Wood, K. V. (2008) HaloTag: a novel protein labeling technology for cell imaging and protein analysis. ACS Chem Biol. 3, 373–382

21. Rinis, N., Golden, J. E., Marceau, C. D., Carette, J. E., Van Zandt, M. C., Gilmore, R., and Contessa, J. N. (2018) Editing N-Glycan Site Occupancy with Small-Molecule Oligosaccharyltransferase Inhibitors. Cell Chem Biol. 25, 1231–1241.e4

22. Foulquier, F., Harduin-Lepers, A., Duvet, S., Marchal, I., Mir, A. M., Delannoy, P., Chirat, F., and Cacan, R. (2002) The unfolded protein response in a dolichyl phosphate mannose-deficient Chinese hamster ovary cell line points out the key role of a demannosylation step in the quality-control mechanism of N-glycoproteins. Biochemical Journal. 362, 491

23. Polla, D. L., Edmondson, A. C., Duvet, S., March, M. E., Sousa, A. B., Lehman, A., Niyazov, D., van Dijk, F., Demirdas, S., van Slegtenhorst, M. A., Kievit, A. J. A., Schulz, C., Armstrong, L., Bi, X., Rader, D. J., Izumi, K., Zackai, E. H., de Franco, E., Jorge, P., Huffels, S. C., Hommersom, M., Ellard, S., Lefeber, D. J., Santani, A., Hand, N. J., van Bokhoven, H., He, M., and de Brouwer, A. P. M. (2021) Bi-allelic variants in the ER quality-control mannosidase gene EDEM3 cause a congenital disorder of glycosylation. Am J Hum Genet. 108, 1342–1349

24. Budhraja, R., Joshi, N., Radenkovic, S., Kozicz, T., Morava, E., and Pandey, A. (2024) Dysregulated proteome and N-glycoproteome in ALG1-deficient fibroblasts. Proteomics. 2024, 2400012

25. Baerenfaenger, M., Post, M. A., Langerhorst, P., Huijben, K., Zijlstra, F., Jacobs, J. F. M., Verbeek, M. M., Wessels, H. J. C. T., and Lefeber, D. J. (2023) Glycoproteomics in Cerebrospinal Fluid Reveals Brain-Specific Glycosylation Changes. Int J Mol Sci. 10.3390/IJMS24031937/S1

26. Kretzmer, C., Narasimhan, R. L., Lal, R. D., Balassi, V., Ravellette, J., Kotekar Manjunath, A. K., Koshy, J. J., Viano, M., Torre, S., Zanda, V. M., Kumravat, M., Saldanha, K. M. R., Chandranpillai, H., Nihad, I., Zhong, F., Sun, Y., Gustin, J., Borgschulte, T., Liu, J., and Razafsky, D. (2022) De novo assembly and annotation of the CHOZN® GS-/- genome supports high-throughput genome-scale screening. Biotechnol Bioeng. 119, 3632–3646

27. Hilliard, W., MacDonald, M. L., and Lee, K. H. (2020) Chromosome-scale scaffolds for the Chinese hamster reference genome assembly to facilitate the study of the CHO epigenome. Biotechnol Bioeng. 117, 2331–2339

28. Kaiden, A., Rosenwald, A. G., Cacan, R., Verbert, A., and Krag, S. S. (1998) Transfer of two oligosaccharides to protein in a Chinese hamster ovary cell B211 which utilizes polyprenol for its N-linked glycosylation intermediates. Arch Biochem Biophys. 358, 303–312

29. Stanley, P. (1983) Lectin-resistant CHO cells: selection of new mutant phenotypes. Somatic Cell Genet. 9, 593–608

30. Sallustio, S., and Stanley, P. (1990) Isolation of Chinese hamster ovary ribosomal mutants differentially resistant to ricin, abrin, and modeccin. Journal of Biological Chemistry. 265, 582–588

31. Lu, H., Sathe, A. A., Xing, C., and Lehrman, M. A. (2019) The Lec5 glycosylation mutant links homeobox genes with cholesterol and lipid-linked oligosaccharides. Glycobiology. 29, 106–109

32. Xu, X., Nagarajan, H., Lewis, N. E., Pan, S., Cai, Z., Liu, X., Chen, W., Xie, M., Wang, W., Hammond, S., Andersen, M. R., Neff, N., Passarelli, B., Koh, W., Fan, H. C., Wang, J., Gui, Y., Lee, K. H., Betenbaugh, M. J., Quake, S. R., Famili, I., Palsson, B. O., and Wang, J. (2011) The genomic sequence of the Chinese hamster ovary (CHO)-K1 cell line. Nature Biotechnology 2011 29:8. **29**, 735–741

33. Huhn, S., Chang, M., Kumar, A., Liu, R., Jiang, B., Betenbaugh, M., Lin, H., Nyberg, G., and Du, Z. (2022) Chromosomal instability drives convergent and divergent evolution toward advantageous inherited traits in mammalian CHO bioproduction lineages. iScience. 10.1016/J.ISCI.2022.104074

34. Gründahl, J. E. H., Guan, Z., Rust, S., Reunert, J., Müller, B., Du Chesne, I., Zerres, K., Rudnik-Schöneborn, S., Ortiz-Brüchle, N., Häusler, M. G., Siedlecka, J., Swiezewska, E., Raetz, C. R. H., and Marquardt, T. (2012) Life with too much polyprenol: polyprenol reductase deficiency. Mol Genet Metab. 105, 642–651

35. Wheeler, P. G., Ng, B. G., Sanford, L., Sutton, V. R., Bartholomew, D. W., Pastore, M. T., Bamshad, M. J., Kircher, M., Buckingham, K. J., Nickerson, D. A., Shendure, J., and Freeze, H. H. (2016) SRD5A3-CDG: Expanding the phenotype of a congenital disorder of glycosylation with emphasis on adult onset features. Am J Med Genet A. 170, 3165

36. Morava, E., Schatz, U. A., Torring, P. M., Abbott, M.-A., Baumann, M., Brasch-Andersen, C., Chevalier, N., Dunkhase-Heinl, U., Fleger, M., Haack, T. B., Nelson, S., Potelle, S., Radenkovic, S., Bommer, G. T., Schaftingen, E. Van, and Veiga-Da-Cunha, M. (2021) Impaired glucose-1,6-biphosphate production due to bi-allelic PGM2L1 mutations is associated with a neurodevelopmental disorder. The American Journal of Human Genetics. 108, 1151–1160

37. Corey, E. J., and Schmidt, G. (1979) Useful procedures for the oxidation of alcohols involving pyridinium dichromate in aprotic media. Tetrahedron Lett. 20, 399–402

38. Kale, D., Kikul, F., Phapale, P., Beedgen, L., Thiel, C., and Brügger, B. (2023) Quantification of Dolichyl Phosphates Using Phosphate Methylation and Reverse-Phase Liquid Chromatography-High Resolution Mass Spectrometry. Anal Chem. 95, 3210–3217

39. Dewulf, J. P., Gerin, I., Rider, M. H., Veiga-Da-Cunha, M., Van Schaftingen, E., and Bommer, G. T. (2019) The synthesis of branched-chain fatty acids is limited by enzymatic decarboxylation of ethyl- and methylmalonyl-CoA. Biochem J. 476, 2427–2447

40. De Coster, W., and Rademakers, R. (2023) NanoPack2: population-scale evaluation of long-read sequencing data. Bioinformatics. 10.1093/BIOINFORMATICS/BTAD311

41. Kolmogorov, M., Yuan, J., Lin, Y., and Pevzner, P. A. (2019) Assembly of long, error-prone reads using repeat graphs. Nat Biotechnol. 37, 540–546

42. Vaser, R., Sović, I., Nagarajan, N., and Šikić, M. (2017) Fast and accurate de novo genome assembly from long uncorrected reads. Genome Res. 27, 737–746

43. Mikheenko, A., Prjibelski, A., Saveliev, V., Antipov, D., and Gurevich, A. (2018) Versatile genome assembly evaluation with QUAST-LG. Bioinformatics. 34, i142–i150

44. Li, H. (2021) New strategies to improve minimap2 alignment accuracy. Bioinformatics. 37, 4572–4574

45. Pedersen, B. S., and Quinlan, A. R. (2018) Mosdepth: quick coverage calculation for genomes and exomes. Bioinformatics. 34, 867–868

46. Thorvaldsdóttir, H., Robinson, J. T., and Mesirov, J. P. (2013) Integrative Genomics Viewer (IGV): high-performance genomics data visualization and exploration. Brief Bioinform. 14, 178–192

47. Nattestad, M., Aboukhalil, R., Chin, C. S., and Schatz, M. C. (2021) Ribbon: intuitive visualization for complex genomic variation. Bioinformatics. 37, 413–415

48. Kentache, T., Thabault, L., Peracchi, A., Frédérick, R., Bommer, G. T., and van Schaftingen, E. (2020) The putative Escherichia coli dehydrogenase YjhC metabolises two dehydrated forms of N-acetylneuraminate produced by some sialidases. Biosci Rep. 10.1042/BSR20200927

49. Okonechnikov, K., Golosova, O., Fursov, M., Varlamov, A., Vaskin, Y., Efremov, I., German Grehov, O. G., Kandrov, D., Rasputin, K., Syabro, M., and Tleukenov, T. (2012) Unipro UGENE: a unified bioinformatics toolkit. Bioinformatics. 28, 1166–1167

