## Supplementary information for "The N-glycosylation defect in Lec5 and Lec9 CHO cells is caused by absence of the DHRSX gene"

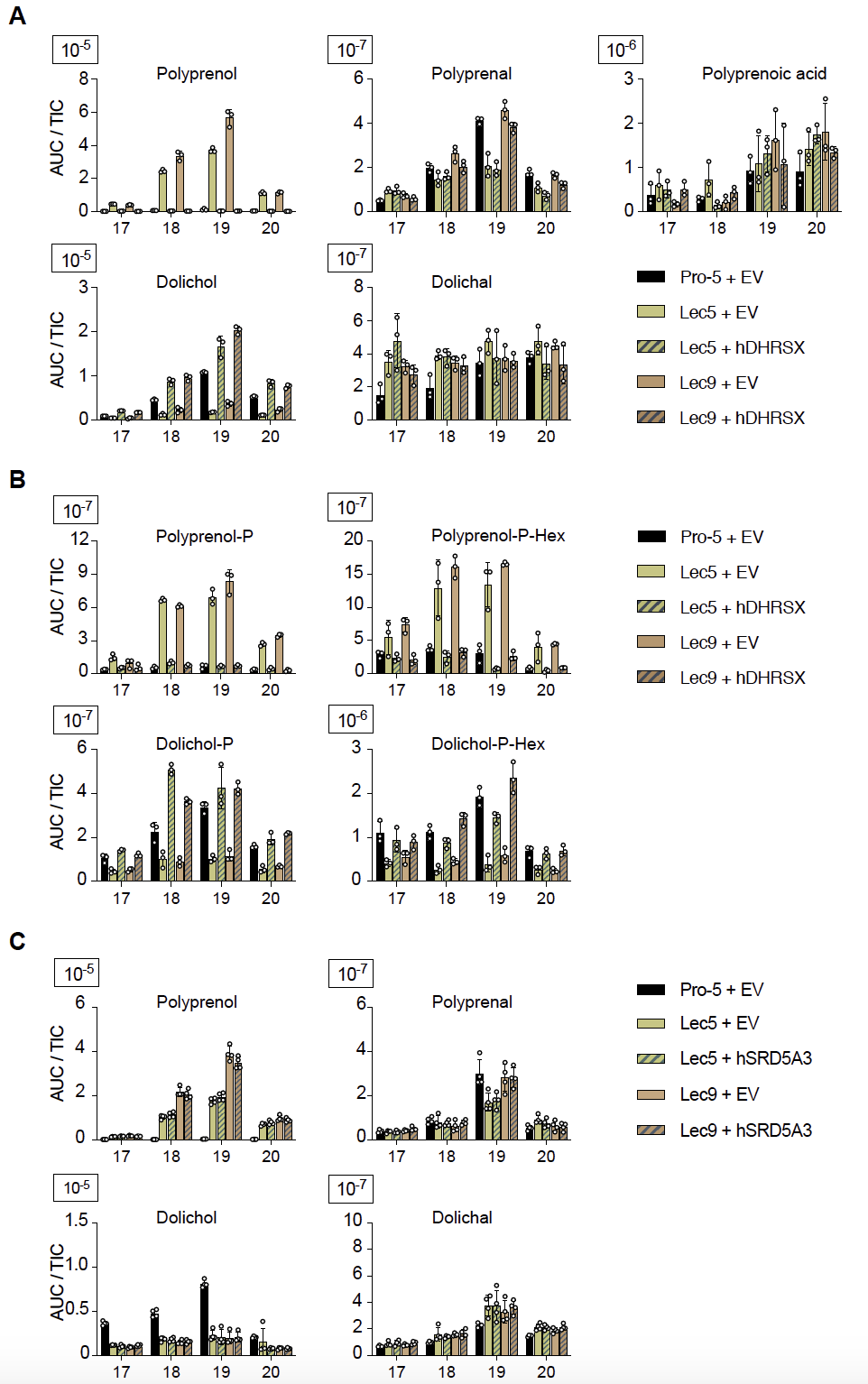

**Figure S1: Additional data complementing Fig. 1 showing isoprenoids with 17-20 species**

(A) Isoprenoid species with 17-20 isoprenyl units in CHO Pro-5, Lec5, Lec9 cells and their respective complementations by human DHRSX. Data are TIC-normalized AUC (mean ± SEM, n=3).

(B) Isoprenoid species with 17-20 isoprenyl units in CHO Pro-5, Lec5, Lec9 cells and their respective complementations by human SRD5A3. Data are TIC-normalized AUC (mean ± SEM, n=3).

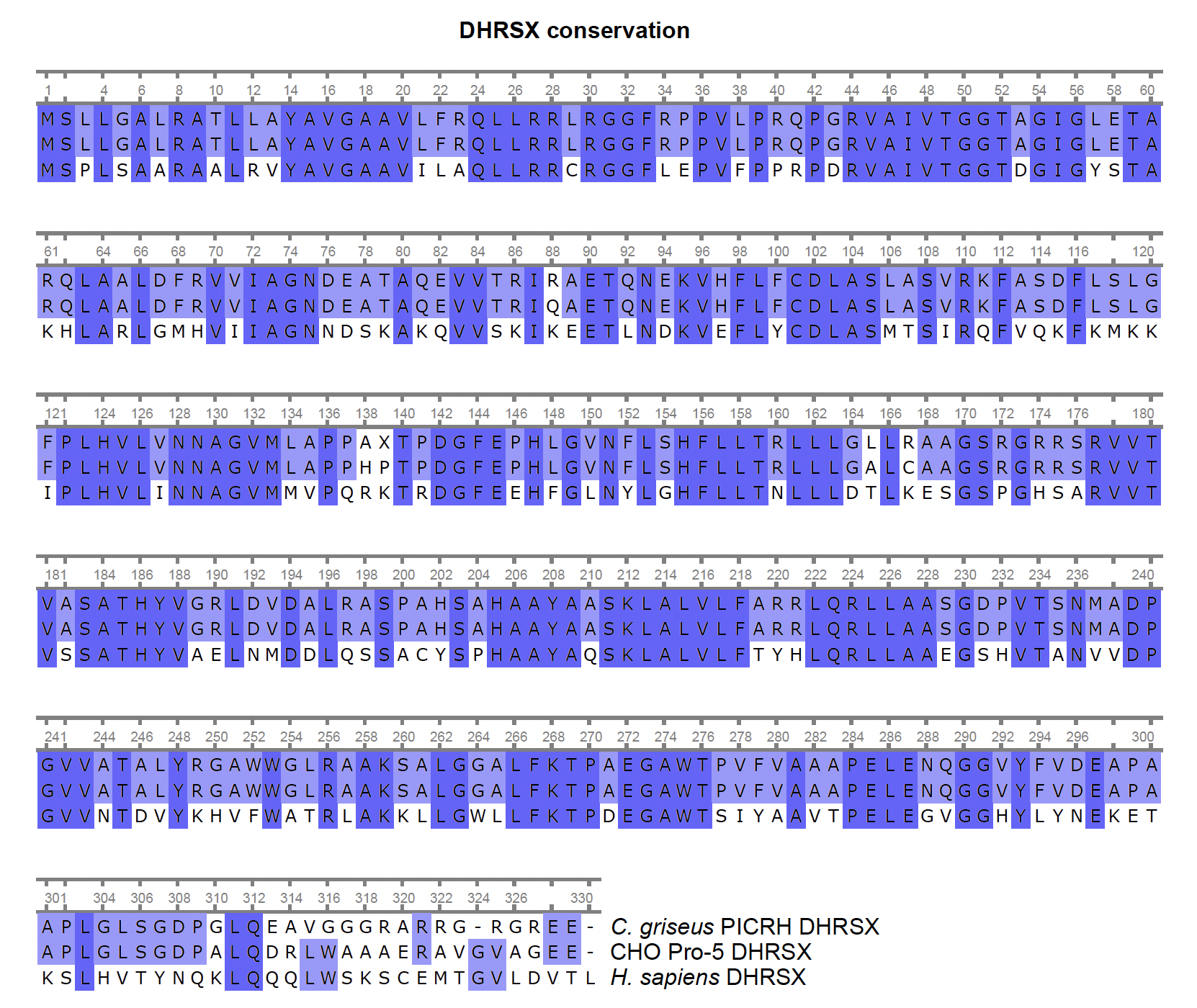

**Figure S2. DHRSX conservation**

Conservation of the DHRSX protein sequence in the *C. griseus* PICRH-1.0 assembly (GenBank: GCA_003668045.2), the sequence detected in our *de novo* assembly from Pro-5 CHO cells, and *H. sapiens* (Uniprot: Q8N5I4). Alignment performed using the ClustalW plugin for Unipro Ugene 49.1 (46).

**Table S1. Non-normalized proteomics data, complementing Figure 4.**

**Table S2. Pro-5, Lec5 and Lec9 *de* novo assembly metrics.** N50: Length of the shortest contig at 50% of assembly length (i.e. 50% of assembly is in contigs of this length or longer); N90: Length of the shortest contig at 90% of assembly length (i.e. 90% of assembly is in contigs of this length or longer); L50: The minimum number of contigs that, combined, make up 50% of the assembly length.

**Table S3. Mean read depth, per contig, of long-reads from Pro-5, Lec5 and Lec9 cells mapped to the *C. griseus* PICRH-1.0 assembly reference genome.**

**Table S4. Mean read depth, per contig, of the genomes derived from Pro-5, Lec5 and Lec9 Flye *de novo* assemblies, mapped to the PICRH-1.0 assembly reference genome**

**Table S5. Theoretical *m*/*z* values of [M + NH_4_^+^] ions of polyprenal, dolichal, polyprenol, dolichol, dolichol M+2, dimethylated polyprenol-phosphate, and dimethylated dolichol phosphate from species with 17 to 21 isoprene units**. The theoretical *m*/*z* values of polyprenoic acid, polyprenol-P-hexose and dolichol-P-hexose correspond to the [M - H^+^] ions.
