## Supplementary material for "The N-glycosylation defect in Lec5 and Lec9 CHO cells is caused by absence of the DHRSX gene": Key resources table

| **Reagent or Resource** | **Source** | **Identifier** |
| --- | --- | --- |
| **Antibodies/lectins** |  |  |
| DHRSX | Sigma-Aldrich | HPA003035 |
| β-tubulin | Thermo Fisher Scientific | MA516308 |
| HPR-linked anti-rabbit IgG | Bioké | 7074S |
| HPR-linked anti-mouse IgG | Bioké | 7076S |
| HaloTag® | Promega | G9211 |
| **Chemicals, reagents and equipment** | | |
| Bovine Serum Albumin (BSA) Fraction V | Sigma | 10735086001 |
| Penicillin/Streptomycin Solution (100x) | CAPRICORN Scientific | CP21-4278 |
| Fetal Bovine Serum | Dutcher | S1810 |
| Nitrocellulose membrane | Invitrogen | LC2000 |
| Fetal Bovine Serum | Sigma-Aldrich | 98022 |
| Lipofectamine 3000 | Thermo Fisher Scientific | 15292465 |
| DMEM/F-12 | Gibco | 21041-025 |
| DMEM/F-12, HEPES | Gibco | 31330038 |
| Cytiva HyClone™ Fetal Clone III Serum | Cytiva | 12319862 |
| Triton™ X-100 | Sigma-Aldrich | T8787 |
| beta-mercaptoethanol | Sigma-Aldrich | M3148 |
| Pageruler Plus Prestained Protein Ladder | Thermo Fisher Scientific | 11832124 |
| DMSO | Sigma-Aldrich | 317275 |
| Para-formaldehyde | Thermo Fisher Scientific | 28908 |
| Mowiol® 4-88 | Sigma-Aldrich | 81381 |
| RIPA buffer | Custom | N/A |
| SuperSignal™ West Pico PLUS Chemiluminescent Substrate | Thermo Fisher Scientific | Cat# 34579 |
| Puromycin Dihydrochloride | Gibco | A1113802 |
| G-418 Geneticin™ Selective Antibiotic | Gibco | 10131035 |
| Tunicamycin | Sigma-Aldrich | SML1287 |
| MG-132 | VWR | 474787 |
| Pyridinium Dichromate | Merck life science BV | 214698-100G |
| PBS | Thermo Fisher Scientific | 10010023 |
| TBS 10X | Euromedex | ET220 |
| Versene | Gibco | 15040033 |
| Methanol | Thermo Fisher Scientific | 10606652 |
| Methanol (LC-MS grade) | Biosolve | 136878 |
| Chloroform | VWR | 22711290 |
| Acetic Acid,99.8% for analysis | Thermo scientific | 222140010 |
| Dichloromethane | Merck | 1.06050.1000 |
| Chloroform (LC-MS grade) | Biosolve | 34806 |
| trimethylsilyl diazomethane (TMSD) | Sigma-Aldrich | 362832 |
| NAD^+^ free acid grade II | ROCHE | 10127990001 |
| NADH disodium salt | ROCHE | 10128023001 |
| NADP^+^ sodium salt hydrate | Sigma-Aldrich | N0505-1G |
| NADPH tetrasodium salt | ROCHE | 10102824001 |
| Dolichol | Aventi polar lipids | 9002000 |
| Polyprenol | Aventi polar lipids | 9002100 |
| Dolichal | This study | N/A |
| Polyprenal | Aventi polar lipids (discontinued) | 9002200 |
| Phosphatidylcholine | Sigma-Aldrich | P-0378 |
| Phosphatidylethanolamine | Sigma-Aldrich | P-0503 |
| Carbenicillin | VWR | J67159.AD |
| Gelred nucleic acid stain | Sigma-Aldrich | SCT123 |
| Agilent 6546 ion funnel mass spectrometer | Agilent | N/A |
| Agilent 1290 HPLC System | Agilent | N/A |
| Accucore C30 150 x 2.1 mm column | Thermo Fisher Scientific | 27826-152130 |
| EASY-Spray 0.075 x 250 mm | Thermo Fisher Scientific | ES902 |
| Trap-column | Thermo Fisher Scientific | Acclaim PepMap100 |
| Isopropopanol (MS-grade) | Biosolve | 162678 |
| Acetonitrile | Biosolve | 12078 |
| Ammonium formate | Biosolve | 19878 |
| Formic acid | Biosolve | 232478 |
| iBlot® 2 NC mini Stacks | Invitrogen | IB23002 |
| ECL WB substrate | Thermo Fisher Scientific | PIER32106 |
| DNAse I | Sigma Aldrich | 10104159001 |
| Trypsin | Gibco | 25300096 |
| Tris-buffered saline | Bioké | 124985 |
| IPTG | Thermo Fisher Scientific | 15529019 |
| Bolt® 4-12% Bis-Tris Plus Gels, 12-well | Thermo Fisher Scientific | 15324604 |
| Bolt Transfer Buffer (20X)-1 L | Thermo Fisher Scientific | 15256066 |
| 20X Bolt® MES SDS Running Buffer (500 mL) | Thermo Fisher Scientific | 13266499 |
| LDS Sample buffer | Thermo Fisher Scientific | 11549166 |
| cOmplete(TM), Mini, EDTA-free Protease | Thermo Fisher Scientific | 11836170001 |
| Trypsin | Sigma-Aldrich | T8003-500MG |
| Trypsin-EDTA 1X in PBS w/o Calcium w/o Magnesium | Dominique Dutscher | L0940-100 |
| Tween-20 | Sigma-Aldrich | P1379 |
| N-Glycosidase F | Roche | 11365193001 |
| 6X DNA Loading Dye | Thermo Fisher Scientific | R0611 |
| Polybrene | Merck life science N.V (ex Sigma Aldrich) | TR-1003 |
| Phusion™ DNA polymerase | Thermo Fisher Scientific | F630S |
| T4 DNA ligase, 5 U/µL | Thermo Fisher Scientific | EL0014 |
| Trizma® base | Merck Life Science BV (ex Sigma Aldrich) | T1503-100g |
| HEPES | Merck Life Science BV (ex Sigma Aldrich) | H3375-500G |
| Antipaine | Merck Life Science B.V. (Sigma) | A6191 |
| Leupeptine | Merck Life Science B.V. (Sigma) | L2884 |
| PMSF | Merck Life Science B.V. (Sigma) | 10837091001 |
| Phosphate buffered saline PBS TABLETS | Millipore | 2810306 |
| PNGase F | NEB | P0704S |
| S-Trap™ micro units | Protifi LLC | N/A |
| HisTrap-HP 1 ml | GE Healthcare | GE29-0510-21 |
| Imidazole | Merck Life Science BV (ex Sigma Aldrich) | 56750-500G |
| Water ULC/MS - CC/SFC | Biosolve | 232141 |
| Sequencing Grade Modified Trypsin (1x100ug) | Promega | V5117 |
| Lys-C endopeptidase | Sopachem NV | 125-02543 |
| Phosphoric acid | Merck Life Science BV (ex Sigma Aldrich) | 695017-100ML |
| Sodium dodecyl sulfate SDS | Merck Life Science BV (ex Sigma Aldrich) | 62862-1KG |
| **Commercial Assays/Kits** | | |
| Micro BCA™ Protein Assay Kit | Thermo Fisher Scientific | Cat# 23235 |
| Ligation Sequencing Kit XL V14 | ONT | SQL-LSK114-XL |
| HS Large Fragment 50kb Kit | Agilent Technologies | DNF-464-33 |
| Pierce^TM^ Quantitative Peptide Assay | Thermo Fisher Scientific | 23275 |
| Monarch^®^ HMW DNA Extraction Kit | NEB | T3050L |
| **Experimental Models: Cell Lines/Strains** | | |
| HEK293T | Gift from Reid Gilmore, Uni. Of Mass. | This paper |
| BL21(DE3) *E. Coli* | Thermo Fisher Scientific | 10749734 |
| XL1-Blue Competent Cells *E. coli* | Agilent | 200249 |
| Oligonucleotides |  |  |
| SRD5A3_rev_bsrGI  ATACAATGTACACTAGAAGGCACAGTCGAGGC | This paper | N/A |
| SRD5A3_sense_BglII  TTATATAGATCTCCACCATGGCTCCCTGGGCGGAG | This paper | N/A |
| hDHRSX_rev_bsrGI  ATACCATGTACATCACAGGGTCACATCAAGGAC | This paper | N/A |
| hDHRSX_ sense_BglII  TTATATAGATCTCCACCATGTCGCCATTGTCTGCGG | This paper | N/A |
| **Plasmids & Vectors** |  |  |
| pET15b DHRSX-N-His NM_145177.3 | This paper | N/A |
| pET15b chDHRSX-N-His | This paper | N/A |
| pcDNA3.1(+) SRD5A3-N-His NM_024592.5 | This paper | N/A |
| pcDNA3-ER-Halo3N | Rinis *et al.* 2018 | N/A |
| pUB83 hSRD5A3-N-His | This paper | N/A |
| pUB83 hDHRSX-N-His | This paper | N/A |
| **Software and Algorithms** |  |  |
| ImageJ 2.14.0 | NIH | <https://www.imagej.net/ij/> |
| Fiji | NIH | doi:10.1038/nmeth.2019 |
| Adobe Illustrator | Adobe | [https://www.adobe.com](https://www.adobe.com/) |
| Interactive Genomics Viewer | Broad Institute | <https://software.broadinstitute.org/software/igv/> |
| Ugene | Unipro | <https://www.ugene.net/> |
| Graphpad Prism 10.1 | Dotmatics | [https://www.graphpad.com](https://www.graphpad.com/) |
| Mass Hunter Quantitative Analysis | Agilent | https://www.agilent.com/en/product/software-informatics/mass-spectrometry-software/data-analysis/quantitative-analysis |
| Mass Hunter Qualitative Analysis | Agilent | https://www.agilent.com/en/product/software-informatics/mass-spectrometry-software/data-analysis/quantitative-analysis |
| Proteome Discoverer 2.5 SP1 | Thermo Fisher Scientific | https://www.thermofisher.com/be/en/home/industrial/mass-spectrometry/liquid-chromatography-mass-spectrometry-lc-ms/lc-ms-software/multi-omics-data-analysis |
| MiniKNOW Core 5.4.3 | ONT | https://nanoporetech.com/news/news-introducing-new-minknow-app |
| NanoPlot 1.41.0 | PyPI | https://pypi.org/project/NanoPlot/ |
| Flye assembler 2.9.2 | Bioconda | https://anaconda.org/bioconda/flye |
| Mosdepth 0.3.5 | Bioconda | https://anaconda.org/bioconda/mosdepth |
| racon 1.5.0 | ILRI Research Computing | https://hpc.ilri.cgiar.org/racon-software |
| QUAST 5.2.0 | QUAST | https://quast.sourceforge.net/docs/manual.html |
| Integrative Genomics Viewer (IGV) 2.17.1 | Integrative Genomics Viewer | https://igv.org/ |
| Protparam tool | expasy | https://web.expasy.org/protparam/ |
