## Supplementary material for "The N-glycosylation defect in Lec5 and Lec9 CHO cells is caused by absence of the DHRSX gene": Table S5

**Table S4:**

**Theoretical *m*/*z* values of [M + NH_4_^+^] ions of polyprenal, dolichal, polyprenol, dolichol, dolichol M+2, dimethylated polyprenol-phosphate, and dimethylated dolichol phosphate from species with 17 to 21 isoprene units. The theoretical *m*/*z* values of polyprenoic acid, polyprenol-P-hexose and dolichol-P-hexose correspond to the [M - H^+^] ions.**

|  | Number of isoprene units | 17 | 18 | 19 | 20 | 21 |
| --- | --- | --- | --- | --- | --- | --- |
| [M + NH_4_^+^] | Polyprenal | 1191.093 | 1259.1556 | 1327.2182 | 1395.2808 | 1463.3434 |
|  | Polyprenol / Dolichal | 1193.1086 | 1261.1712 | 1329.2338 | 1397.2964 | 1465.359 |
|  | Dolichol | 1195.1224 | 1263.1868 | 1331.2494 | 1399.312 | 1467.3746 |
|  | Dolichol M + 2 | 1197.1291 | 1265.1935 | 1333.2561 | 1401.3187 | 1469.3813 |
|  | Dimethylated Dolichol-Phosphate | 1303.12188 | 1371.18448 | 1439.24708 | 1507.30968 | 1575.37228 |
|  | Dimethylated Polyprenol  -Phosphate | 1301.10623 | 1369.16883 | 1437.23143 | 1505.29403 | 1573.35663 |
| [M - H^+^] | Polyprenoic acid | 1188.0467 | 1256.1093 | 1324.1719 | 1392.2345 | 1460.2971 |
|  | Dolichol-P-hexose | 1418.1023 | 1486.1648 | 1554.2274 | 1622.2901 | 1690.3527 |
|  | Polyprenol-P-hexose | 1416.0866 | 1484.1492 | 1552.2118 | 1620.2744 | 1688.3371 |
